# Landscape and regulation of mRNA translation in the early *C. elegans* embryo

**DOI:** 10.1101/2024.12.13.628416

**Authors:** Yash Shukla, Vighnesh Ghatpande, Cindy F. Hu, Daniel J. Dickinson, Can Cenik

## Abstract

Animal embryos rely on regulated translation of maternally deposited mRNAs to drive early development. Using low-input ribosome profiling combined with RNA sequencing on precisely staged embryos, we measured mRNA translation during the first four cell cycles of *C. elegans* development. We uncovered stage-specific patterns of developmentally coordinated translational regulation. We confirmed that mRNA localization correlates with translational efficiency, though initial translational repression in germline precursors occurs before P-granule association. Our analysis suggests that the RNA-binding protein OMA-1 represses the translation of its target mRNAs in a stage-specific manner, while indirectly promoting the translational efficiency of other transcripts. These findings illuminate how post-transcriptional mechanisms shape the embryonic proteome to direct cell differentiation, with implications for understanding similar regulation across species where maternal factors guide early development.

## 1. Introduction

During animal development, a single-cell zygote undergoes cleavage to produce founder blastomeres that subsequently form specialized organs and tissues. The early embryo must precisely regulate protein expression both spatially and temporally, despite being transcriptionally quiescent to varying degrees^1^. To overcome this challenge, embryos rely on post-transcriptional regulation of maternally deposited proteins and mRNAs^2^. Post-transcriptional regulation is particularly critical prior to the maternal-to-zygotic transition, when the embryo begins to rely on newly generated transcripts from its own genome^3^. However, post-transcriptional regulation continues to play important roles beyond the maternal-to-zygotic transition in many species, as maternal mRNAs influence development even after zygotic genome activation^4,5^.

In *C. elegans*, the embryonic cell lineage is invariant, with cell fates becoming irreversibly determined beginning from the first cleavage^6^. Notably, this irreversible cell differentiation occurs in the absence of new mRNA synthesis^7,8^. This early specification raises the intriguing question of how cell fates are determined before the activation of the zygotic genome. In most other organisms, cell fate determination relies on both transcriptional and translational regulation^2^ ; thus, *C. elegans* offers a unique opportunity to study translational control in isolation during early development. Recent work has further demonstrated that maternal ribosomes alone are sufficient for embryonic development in *C. elegans*^9^, suggesting that the translation machinery established in the oocyte is capable of sustaining early developmental processes.

Although single-cell RNA-seq studies have revealed maternal transcript abundance in individual cells within early *C. elegans* embryos (1-16 cells)^8,10–12^, our understanding of spatiotemporal control of mRNA translation remains limited. Genetic and imaging studies identified a few specific instances of translational regulation, including regional control of POS-1 translation^13–16^ and regulation of other maternal transcripts ^17–20^, but these gene-by-gene studies do not reveal the global landscape of translational regulation in early embryos. Bulk ribosome profiling studies in *C. elegans* embryos have provided insights into translational regulation during embryogenesis^21,22^, but these studies lacked temporal resolution due to the challenge of collecting staged embryo populations, and would average out any stage-specific translational regulation. In this work, we measured mRNA translation in precisely staged *C. elegans* embryos using a recent innovation, ribosome profiling via isotachophoresis (Ribo-ITP)^23^. Ribo-ITP employs a custom designed microfluidic chip that concentrates and size-selects ribosome-protected RNA fragments using on-chip isotachophoresis^24^. This approach enables highly efficient recovery of ribosome footprints from minimal input material. Ribo-ITP has allowed extraction of ribosome footprints from single human cells and single mouse embryos, representing a ∼10,000-fold increase in sensitivity compared to earlier bulk ribosome footprinting approaches^25,26^. Adapting Ribo-ITP to *C. elegans* embryos allowed us to overcome the challenge of rapid and asynchronous embryonic development, which precludes large-scale collections of staged embryos. Here, we applied Ribo-ITP and low-input RNA-seq to analyze the first three cell divisions of *C. elegans* embryogenesis, revealing transcriptome-wide changes in translation of mRNA at the one-, two-, four-, and eight-cell stages of *C. elegans* embryos.

Translational landscapes in early embryos are significantly influenced by maternally deposited RNA-binding proteins^27^. Many of these proteins, when mutated, cause maternal-effect lethality and/or alter blastomere cell fates^28–31^. By analyzing translational changes in an RNA-binding protein mutant we identified stage-specific regulation of hundreds of transcripts by OMA-1^31^, a translational repressor implicated in oocyte development^32–34^ and early embryogenesis^13,18,35,36^. These findings expand our understanding of OMA-1’s regulatory targets during early development in *C. elegans* and contribute to our knowledge of the influence of maternal RNA-binding proteins on stage-specific translation during early embryogenesis.

## 2. Results

### 2.1. Low-input RNA-seq and Ribosome Profiling

We focused on the first three cell divisions of *C. elegans* embryogenesis, a period when the progenitors of all three germ layers are established. For each stage, we hand-dissected and flash-froze pools of nine precisely staged embryos for ribosome profiling (Ribo-ITP) or RNA-seq experiments (Figure 1A). Although we were able to obtain ribosome footprints from single embryos in pilot experiments, we found that using pools of nine embryos provided superior footprinting depth compared to using one embryo.

**Figure 1:**
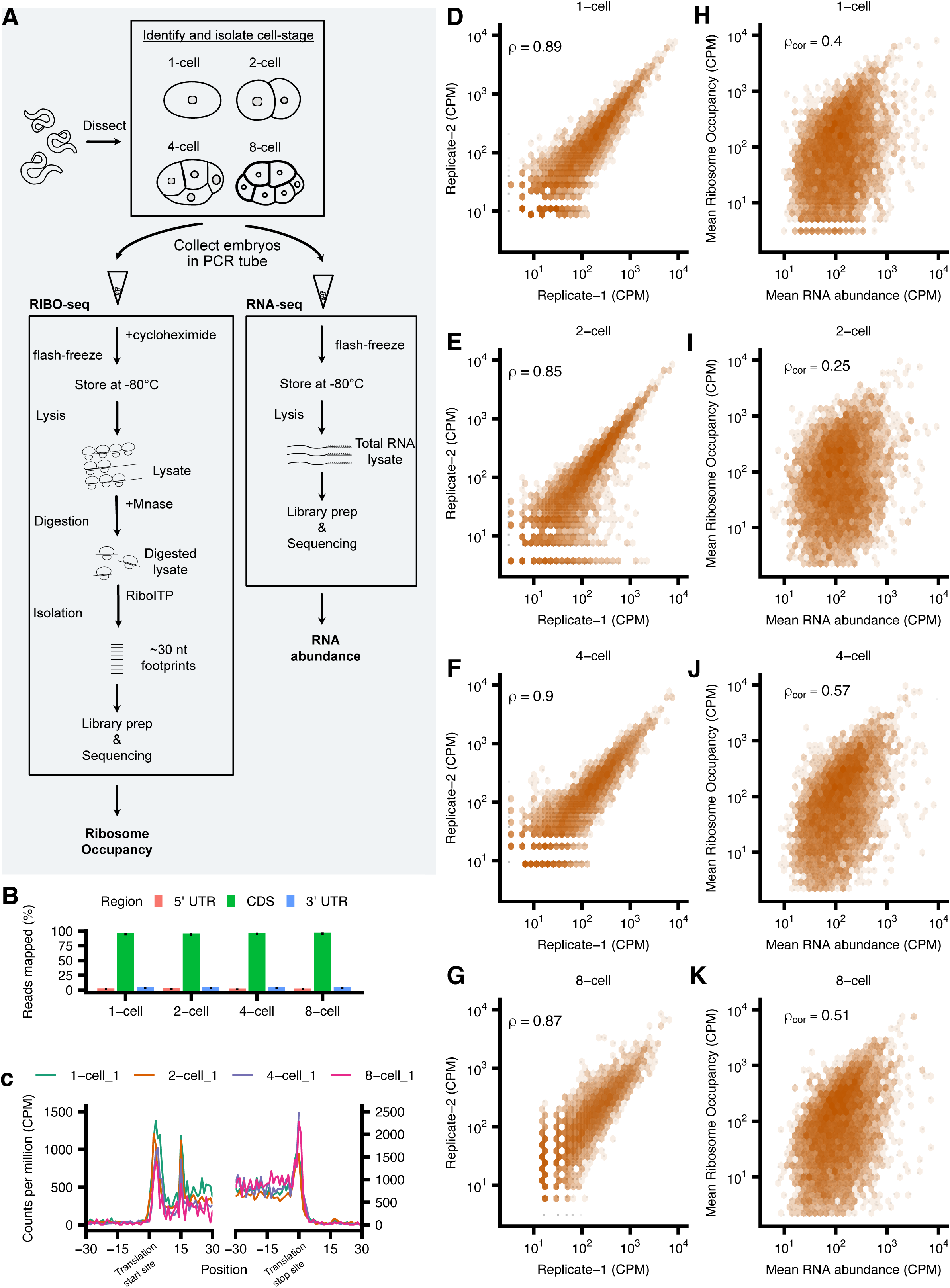
Low-input RNA-seq and Ribosome Profiling. **(A)** Adult *C. elegans* were dissected to obtain embryos at 1-cell, 2-cell, 4-cell, and 8-cell stages, which were then subjected to ribosome profiling (Ribo-ITP) and RNA-seq analysis. **(B)** Mapping of ribosome profiling reads to different genomic features (CDS, 5’ UTR, 3’ UTR) is presented for each stage. Error bars indicate the standard deviation. **(C)** Ribosome occupancy around the translation start and stop sites in a representative 1-cell, 2-cell,4-cell and 8-cell staged embryo. Translation start (or stop) sites are denoted by the position 0. Aggregated read counts (*y* axis) relative to the start (or stop) sites are plotted after A-site correction **(D-G)** Hexbin plot illustrating pairwise correlations of Ribo-seq data between representative replicates across developmental stages. Each axis displays log-scaled counts per million (CPM) of detected genes. The Spearman correlation is indicated in the left corner of the plots. Darker orange color indicates high density of points. **(H-K)** Pairwise correlation between ribosome occupancy and RNA abundance at the four stages of early *C. elegans* embryo development is presented. The mean cpm of ribosome occupancy is plot against the mean cpm of RNA abundance at each stage. p_cor_ is the corrected spearman correlation based on the reliability r (RNA) and r (Ribo) which are the replicate-to-replicate correlation (methods).

For each developmental stage, we obtained at least two high-quality Ribo-ITP replicates, which showed reads predominantly mapping to the coding sequence (CDS) region (Figure 1B) and characteristic enrichments at the start and stop sites (Figure 1C). Furthermore, these CDS-mapped reads exhibited the expected footprint size of approximately 30 nucleotides, typical of ribosome profiling experiments using MNase digestion (Figure S1A). After filtering and quality control, we obtained 4,905 genes that showed robust ribosome occupancy reads(methods). We observed strong replicate-to-replicate correlation between the number of ribosome footprints mapping to coding regions (Spearman rank correlation *ρ >* 0.85 P < 2 × 10^−16^), supporting the method’s reproducibility (Figure 1D-G and Figure S1B). Clustering analysis, using multidimensional scaling, revealed global similarity of translational profiles obtained in replicate experiments, with the exception of one outlier (Figure S1C). Taken together, these metrics indicate technical robustness and reproducibility of our Ribo-ITP measurements.

To estimate how efficiently the mRNA of a given gene is being translated, its ribosome occupancy needs to be normalized to its mRNA abundance, which we quantified using RNA-seq experiments on the same developmental stages as those analyzed by Ribo-ITP. We observed that variability between replicates increased as development progressed, with the mean correlation between replicates decreasing from *ρ* = 0.92 at the 1-cell stage to *ρ* = 0.73 at the 8-cell stage (Figure S1D-E). We attribute this increase in variability to three factors: the stochasticity of gene expression^37^, the increasing complexity of later embryos (with multiple cell types present), and possible PCR duplication during RNAseq library preparation. Despite these issues, in our 8-cell RNAseq dataset we robustly detected genes known to be expressed at the 8-cell stage in *C. elegans*^8,38,39^, verifying the accuracy of our RNA-seq data for capturing developmental progression (Figure S2A). Having collected the two key metrics – RNA abundance and ribosome occupancy – we are now equipped to answer how translation of maternal mRNA changes globally.

### 2.2. Stage-specific patterns of translational regulation in early embryogenesis

The strength of the correlation between mRNA abundance and ribosome occupancy can indicate the extent to which translation is regulated. If there were no translational regulation – that is, if the amount of mRNA translation were governed solely by mRNA abundance – then mRNA abundance and ribosome occupancy should correlate strongly. Strikingly, however, we observed a weak correlation between mRNA abundance and ribosome occupancy, indicating widespread regulation of translation (Figure 1H-K). Furthermore, the correlation between ribosome occupancy and RNA-seq exhibited dynamic changes across early embryonic stages. Starting with a moderate correlation at the 1-cell stage (ρ = 0.361, p < 2e-16; ρ_cor_ = 0.4; Figure 1H), it decreased to a weak correlation at the 2-cell stage (ρ = 0.220, p < 2e-16; ρ_cor_ = 0.25; Figure 1I). The correlation then strengthened again during the 4-cell stage (ρ = 0.502, p < 2e-16; ρ_cor_ = 0.57; Figure 1J) and 8-cell stage (ρ = 0.417, p < 2e-16; ρ_cor_ = 0.51; Figure 1K). This pattern, particularly the sharp drop at the 2-cell stage followed by recovery at the 4-cell stage, suggests that the relationship between mRNA abundance and ribosome occupancy is developmentally regulated and may reflect stage-specific translational control mechanisms during early embryogenesis.

The observed low correlation between mRNA abundance and ribosome occupancy likely reflects the combined effect of technical variability, stage-specific translational regulation, and intrinsic diversity in mRNA translation efficiency (for example, due to differences in codon usage). We reasoned that previous ribosome profiling experiments from mixed-stage embryos would provide information regarding the relative contribution of the last two variables. Ribosome profiling experiments from mixed-stage datasets from pooled 1-cell to 200-cell stage^40^ showed a strong correlation (Spearman rank correlation = 0.9, p < 2e-16; Figure S2B) between mRNA abundance and ribosome occupancy. This high correlation likely represents a situation where intrinsic differences in mRNA properties (such as sequence features affecting translation efficiency) predominate. In contrast, analyses from pooled 1-cell to 4-cell stage embryos^21^ revealed a more moderate correlation (Spearman rank correlation = 0.58, p < 2e-16; Figure S2C). Our stage-specific correlations (Spearman rank correlations ranging from 0.22 to 0.50) were lower still, suggesting that beyond technical variability and intrinsic diversity in mRNA translation efficiency, there may be substantial translational regulation that varies between developmental stages. Importantly, this observation was not due to measurement error, because when we accounted for variation across biological replicates (see methods), we still observed a mean corrected correlation (ρ_cor_) of 0.43 between ribosome occupancy and RNA-seq across all four early embryonic stages. The progressive decrease in correlation strength—from mixed-stage to pooled early stages to individual cell stages—provides evidence that dynamic, stage-specific translational control mechanisms play an important role during early *C. elegans* embryogenesis, beyond what can be explained by inherent differences in mRNA properties alone.

We next wanted to identify which mRNAs are translationally regulated at each stage. We define “translation efficiency” (TE) in alignment with its common usage in the literature^25,26,41–43^ by normalizing ribosome occupancy to mRNA abundance. TE provides a measure of the relative rate of protein synthesis ^25,44,45^ and is an indicator of active translation for most endogenous transcripts^25^, despite some exceptions in specific instances where ribosome density is decoupled from protein production (e.g., ribosome stalling^46,47^).

To assess developmentally regulated translation of maternal mRNAs, we conducted pairwise differential expression analysis between subsequent stages of development, observing changes in RNA abundance, ribosome occupancy and TE. We detected 179 and 40 mRNAs whose TE increased (FDR < 0.2, log2FC > 1) or decreased (FDR < 0.2, log2FC > −1), respectively, as embryos progressed from the 1-cell to the 2-cell stage (Figure 2A). From the 2-cell to the 4-cell stage we observed 30 and 11 mRNAs whose TE was relatively increased or decreased, respectively (Figure 2B). We did not observe significant changes between the 4-cell and 8-cell stages (Figure 2C), which could reflect both technical and biological factors. The 1-cell to 2-cell comparison represents a transition between two relatively uniform states, while later stages involve multiple distinct cell types, potentially making stage-specific changes more difficult to detect. Additionally, rapid and asynchronous cell divisions occurring from 6 to 8 to 12 cells in this system, and/or mixing of multiple distinct cell types within an 8-cell embryo, would tend to dilute out cell-specific translational changes. Comparing 1-cell to 8-cell stages (Figure S3A) revealed broader translational remodeling, with 421 genes showing increased TE and 296 showing decreased TE (|log_2_FC| > 1, FDR < 0.1). The larger number of significant changes detected when comparing 1-cell to 8-cell stages suggests that many transcripts undergo gradual but consistent changes in TE across several cell cycles, some of which do not reach statistical significance when comparing between neighboring stages (e.g., 4-cell vs. 8-cell). The substantial number of differentially translated transcripts - affecting over 700 genes in total - suggests a major reorganization of the translational landscape during early development. This large-scale translational regulation likely reflects the transition from maternally provided programs to stage-specific translation required for proper cell fate specification.

**Figure 2:**
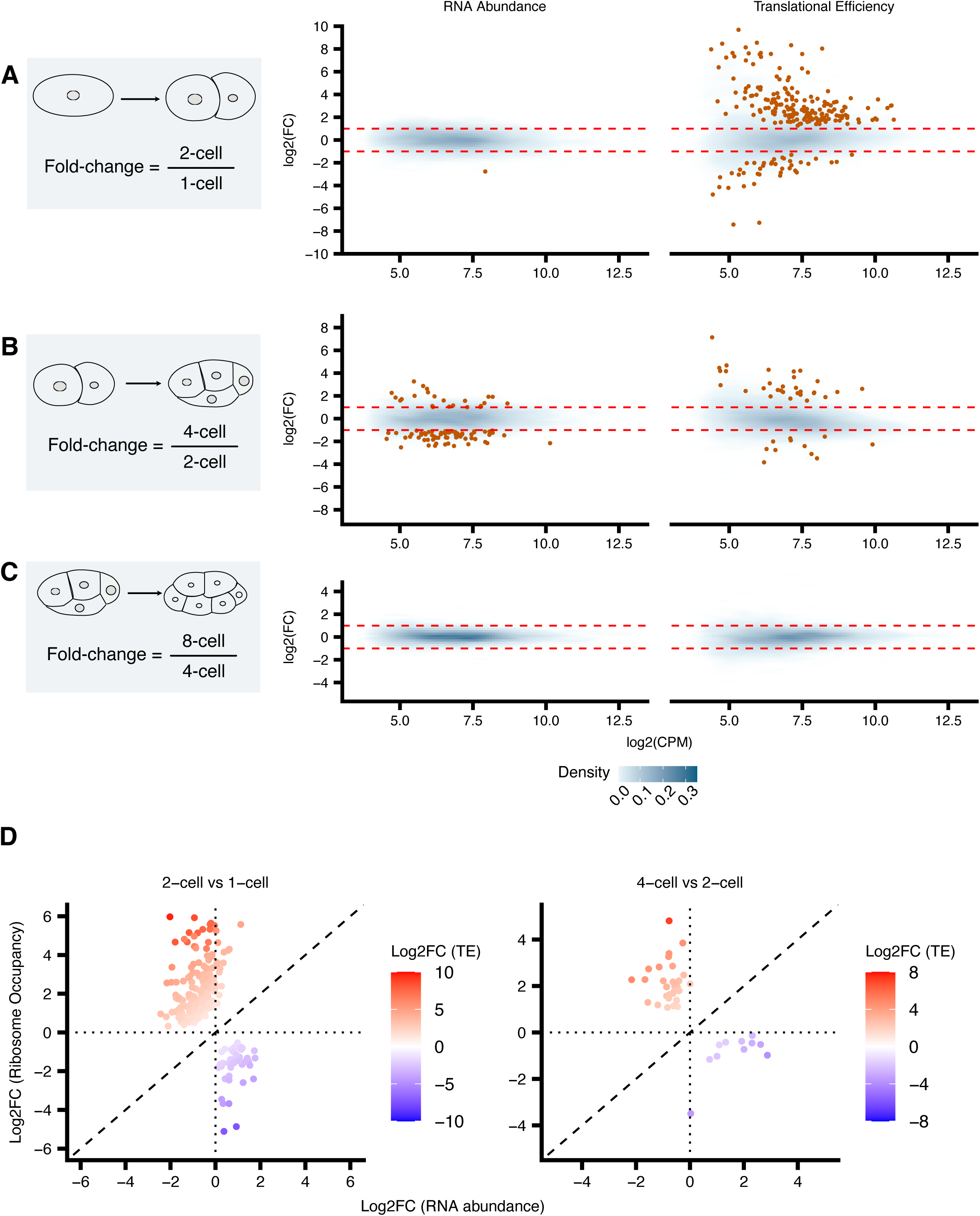
Stage-specific patterns of translational regulation in early embryogenesis. Mean difference plots comparing gene expression between sequential *C. elegans* embryonic stages **(A)**1-cell to 2-cell, **(B)** 2-cell to 4-cell, and **(C)** 4-cell to 8-cell. The log_2_ fold change in RNA abundance and Translational efficiency (y-axis) is plotted against the mean of normalized counts (x-axis). Blue density plot represents the overall distribution of genes with the intensity corresponds to the density of points. The orange points indicate transcripts with significant expression changes (FDR < 0.2 and log_2_FC >1 or < −1). **(D)** Scatter plot comparing the log2 fold change in ribosome occupancy against RNA abundance for genes with significantly altered translational efficiency. The color gradient represents the corresponding translational efficiency for each gene.

We next visualized the extent to which observed changes in TE were due to changes in ribosome occupancy (Figure S3B) vs. changes in mRNA abundance (Figure 2D). For most genes, the change in ribosome occupancy was larger than the change in mRNA abundance. An exception is a cluster of genes whose mRNAs increase 2 to 4-fold in abundance at the 4-cell stage without a corresponding increase in ribosome footprints (Figure 2D, blue points to the right of the origin in the right-hand plot). These apparent increases in mRNA abundance may reflect the compositional nature of RNA-seq and ribosome profiling^48^, whereby degradation of a large number of transcripts^21,49,50^ could artificially inflate the relative abundance of the remaining stable mRNAs. Taken together, we observe a large-scale stage-specific translationally efficiency changes that occur with limited changes in RNA-abundance.

We compared our normalized translational efficiency (see methods) against prior measurements by plotting our measured stage-specific TE values against the TE values reported by Quarto et al. (2021) and Lee et al. (2022). Our data demonstrate concordance with the translational regulation patterns reported by Quarto et al. (2021)^21^ (Spearman rank correlation range from 0.54 to 0.64, p < 2e-16; Figure S3C), while showing weaker similarity to findings in Lee et al. (2022)^22^ (Spearman rank correlation range from 0.36 to 0.42, p < 2e-16; Figure S3D). This alignment with Quarto et al. is expected, as their methodology specifically isolated ribosome profiling data from pooled early embryos, which more closely match our experimental strategy. In contrast, Lee et al. (2022)^22^ analyzed a broader developmental window (1-200 cells) as a single population, likely diluting stage-specific translational signatures that are prominent in our narrowly defined developmental timepoint.

### 2.3. Functional clustering reveals developmentally coordinated translational programs

We next asked whether functional groups of genes undergo coordinated translational regulation across the 1-cell, 2-cell, 4-cell, and 8-cell stages. To compare stage-specific changes in TE, we normalized the TE values to the 1-cell stage for all genes and centered the data by subtracting the mean TE across all stages for each gene (see Methods). This preprocessing step allowed us to highlight developmental stage-specific fluctuations in TE. To analyze coordinated regulation across stages, we employed a group-based approach using k-means clustering to categorize genes based on their TE patterns, resulting in nine distinct clusters (Figure 3A). While a gene-by-gene statistical analysis can identify significantly changed individual genes (Figure 2A-C), this clustering approach enhanced our ability to detect more subtle patterns of coordinated regulation across developmental stages by analyzing groups of genes that underwent similar changes. We observed that most of the changes in TE occurred due to changes in ribosome occupancy, while mRNA abundance was generally stable through these cell stages (Figure 3B). This approach enabled us to identify and characterize groups of genes with similar translational regulation patterns. We observed enrichment of unique biological functions in each cluster (Figure 3C). The clusters with opposing translational trajectories showed particularly distinct functional specializations. For example, in Cluster I, which exhibits progressive translational repression from the 1-cell to 8-cell stage, we observed enrichment of P-granule components and regulators of TOR signaling, perhaps suggesting a transition from oocyte to embryonic development where TOR-mediated nutrient sensing becomes less critical as the embryo shifts to a developmental program less dependent on external nutrient availability. In contrast, Cluster IX, which shows strong translational activation across development, is enriched for transcription factors and chromatin regulators, suggesting that the embryo is preparing for zygotic genome activation by increasing the translation of proteins needed for transcriptional machinery before widespread zygotic transcription begins.

**Figure 3:**
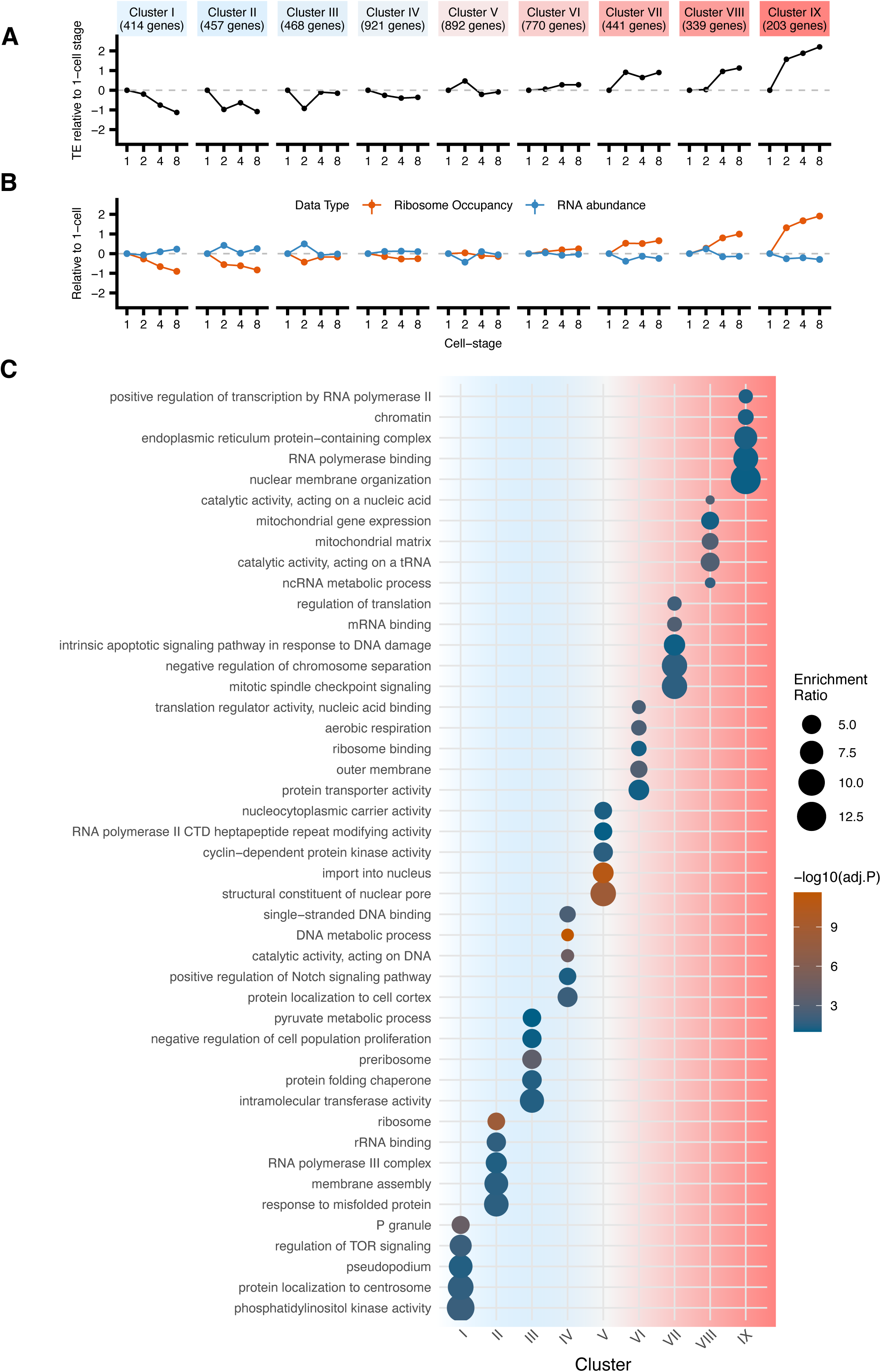
Functional clustering reveals developmentally coordinated translational programs. K-means clustering was performed on normalized TE values for all genes, with data centered by subtracting the mean TE across all stages for each gene. This analysis yielded 9 distinct clusters. **(A)** The line plot displays 1-cell stage subtracted TE values (y-axis) across developmental stages (x-axis). Clusters are color-coded based on broad TE patterns: blue for repression and red of activation. Error bars represent the standard error of TE values for all genes within each cluster. **(B)** Line plot showing the centered-log ratio of ribosome occupancy (orange) and RNA abundance (blue) at developmental stages relative to the 1-cell stage. **(C)** The dot plot visualizes Gene Ontology (GO) enrichment analysis for the identified clusters. The y-axis shows enriched GO terms, while the x-axis represents clusters. Dot size indicates enrichment ratio, and color intensity reflects the adjusted p-value of enrichment.

We next checked whether the patterns we observed were consistent with prior knowledge. Translational regulation has been shown to influence Wnt signaling^18^, E3 ubiquitin ligase-mediated protein degradation^36^, and transcription factor activity^51–53^. We observed that the Wnt protein MOM-2 (cluster VIII) was translationally repressed at the 1-cell and 2-cell stage and increased in TE by ∼10-fold at the 4-cell stage (Figure S4A), consistent with the onset of its protein expression at the 4-cell stage^18^. The transcription factor PAL-1 (cluster VIII; Figure 3A) had a TE 4.5 times lower than that of the average gene at the one-cell stage, with rapid increases at the two- and four-cell stages before leveling off at the 8-cell stage (Figure S4A). Consistent with this, PAL-1 was previously shown to be expressed beginning at the 4-cell stage, although *pal-1* mRNAs are maternally deposited^52,54^. Thus, for these examples, our transcriptome-wide data are consistent with prior knowledge from gene-by-gene studies.

NEG-1 (cluster I) translation was proposed to be regulated by mRNA binding proteins including MEX-5, MEX-3, POS-1, GLD-2, and GLD-3^16^. We observed a progressive decrease ribosome occupancy for cluster I transcripts including *neg-1* from 1-cell to 8-cell stages, with a limited change in overall RNA levels (Figure 3B). Correspondingly, we found reduced TE for *neg-1* at 4-cell and 8-cell stages (Figure S4A), earlier than the previously observed protein degradation at the 10-cell stage^16^.

Finally, the Nanos homolog NOS-2 (cluster VI), whose mRNA is maternally deposited but not translated until the 28-cell stage (and then only in the germline founder cell P4)^13,22,55^, exhibited very low TE at all the stages we analyzed (Figure S4A), consistent with its strong translational repression. Taken together, our genome-wide translational profiling data mirror known regulatory dynamics of key developmental factors, demonstrating the sensitivity and reliability of our approach in capturing translational regulation of maternal mRNAs.

To further validate our findings and explore potentially uncharacterized changes in protein expression, we examined several key developmental proteins that were identified as being translationally regulated in our dataset. We identified the Wnt pathway receptor tyrosine kinase CAM-1 (Cluster IX) as translationally upregulated beginning at the 2-cell stage (Figure S4A), though its protein expression had only been previously observed in later stages^56^. Initially, we detected only punctate localization of an endogenous CAM-1::mNG fusion in live early embryos (Figure S4B-C) that differed from CAM-1’s expected localization at cell-cell contacts^56^. We speculated that newly translated CAM-1::mNG might be difficult to detect via live imaging due to slow fluorescent protein maturation at the standard *C. elegans* culture temperature of 20°C. To overcome this limitation, we fixed embryos and incubated them at 37°C to allow fluorescent protein maturation before imaging (see Methods). This approach revealed previously unobservable CAM-1 protein signals at early stages, specifically at cell-cell contacts beginning at the 4-cell stage (Figure S4C), consistent with its increase in TE in our Ribo-ITP data. We extended this analysis to LAG-1 (Cluster IX), a Notch pathway transcription factor, which was visible in the nucleus from the 1-cell stage in live imaging but only showed an apparent increase in signal intensity from 1-cell to 4-cell stages after fixation and fluorophore maturation (Figure S4D). Interestingly DSH-2 (Cluster VIII), another Wnt pathway component predicted to increase from 2-cell to 4-cell stages, showed detectable signal increases in both live and fixed conditions (Figure S4E), suggesting that perhaps fluorophore maturation can vary depending on the fusion protein. Collectively, these results demonstrate that key components of developmental signaling pathways are translated earlier than previously appreciated, and that Ribo-ITP can identify previously undetected protein expression in early *C. elegans* embryos.

### 2.4. P-granule localization occurs only after translational repression

The distinction between germline and somatic cell lineages is a fundamental aspect of animal development. To explore how translation is regulated during development of the germline lineage, we integrated our TE measurements with existing single-cell RNAseq^8^ and *in situ* hybridization data^40,57^ that revealed the localization of transcripts in specific cells or subcellular regions. Our analysis revealed significantly lower translation in germline-localized transcripts compared to somatic transcripts across early development (Figure 4A; two-cell: −0.896 vs 0.792; four-cell: −1.44 vs 0.912; eight-cell: −1.34 vs 1.16; all stages p < 0.001).

**Figure 4:**
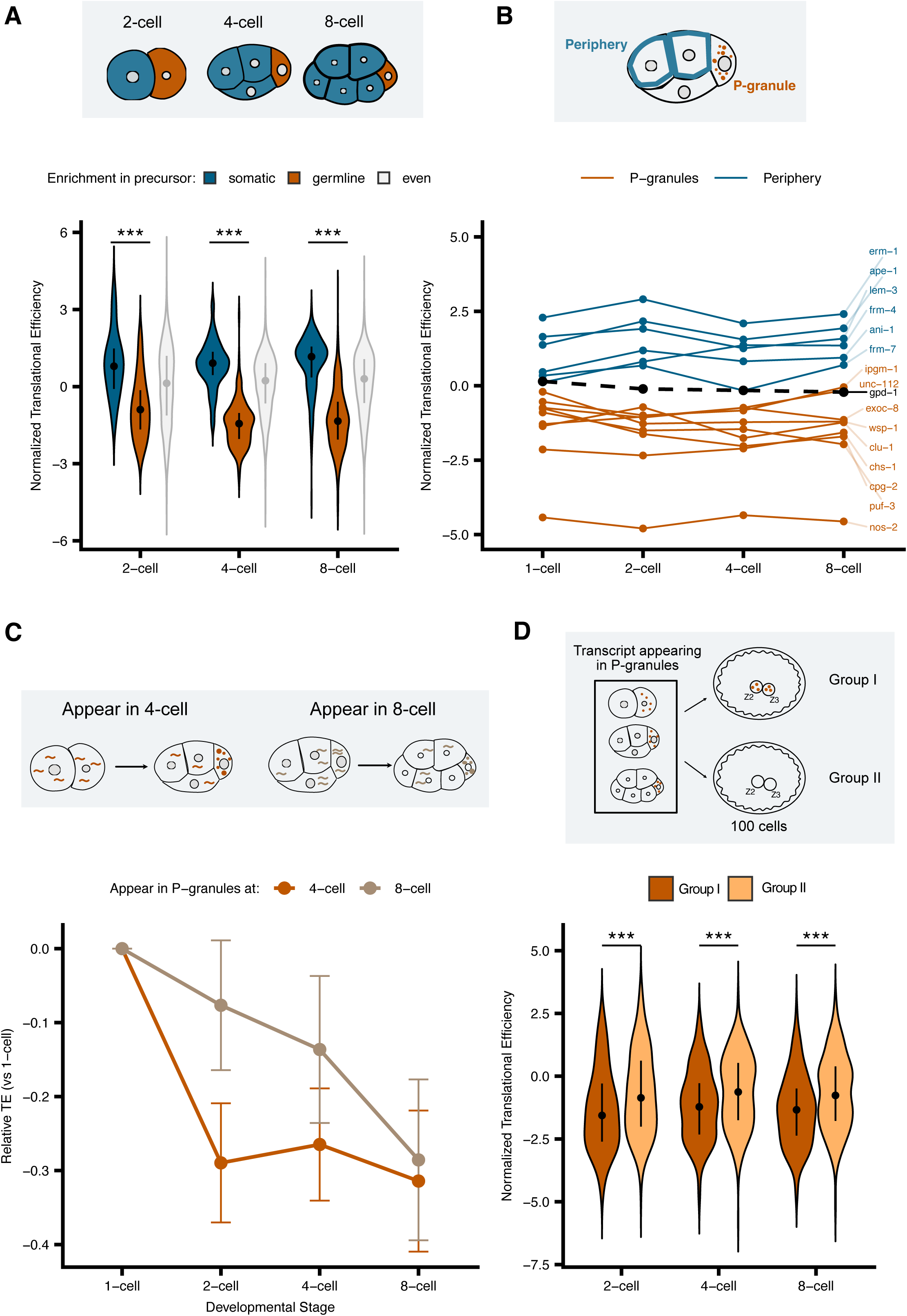
P-granule localization occurs after translational repression. **(A)** Violin plots illustrating the distribution of normalized translational efficiency for transcripts enriched in somatic and germline precursor cells (enrichment data from Tintori et al ^8^) of early *C. elegans* embryos. The width of each violin represents the probability density of efficiency values. Median normalized translational efficiency is indicated by a dot, with error bars showing the first (Q1) and third (Q3) quartiles. **(B)** Line plot showing the mean normalized translational efficiency of transcripts localized in the cell periphery and P-granules (based on smFISH data from Parker et al^57^ and Winkenbach et al ^19^) across embryonic stages from 1-cell to 8-cell. This plot compares the translational efficiency trends of these two transcript populations throughout early embryonic development. **(C)** Translational efficiency relative to 1-cell stage for transcripts that transition into P-granules (P-granule localization identified in Scholl et al ^40^). Orange line shows transcripts that first appear in P-granules at the 4-cell stage (n=81), brown line shows transcripts first appearing at 8-cell stage (n=43). Error bars represent standard error of the mean. **(D)** Translational efficiency of P-granule transcripts based on their maintenance in primordial germ cells (Z2/Z3). Group I transcripts remain associated with P-granules through primordial germ cell development (n=163), while Group II transcripts show transient P-granule association (n=277) (localization identified in Scholl et al). Violin plots show distribution of translational efficiency values, with median and quartiles indicated. Asterisks denote statistical significance between groups (*p < 0.05, **p < 0.01, ***p < 0.001, Wilcoxon rank-sum test).

P granules, which are predominantly found in the germline, have been proposed as sites where translationally repressed transcripts accumulate. When we compared transcripts that were shown to localize to the cell periphery versus in P granules ^19,57^, we observed that P granule localized transcripts showed low TE while cell periphery transcripts showed high TE. To further validate this observation, we analyzed 458 P-granule-localized transcripts from a prior study^40^, which confirmed their globally reduced translational efficiency across development (Figure S5A; median TE at 2-cell: −1.51, 4-cell: −1.20, 8-cell: −1.14). This suggested that translational repression is a specific property of P-granule-associated transcripts.

Since P granules are associated with translational repression, their presence in the germline founder cells could, in principle, explain the reduced translation of germline-enriched transcripts. However, when we examined the overlap between P-granule transcripts (obtained from Scholl et al ^40^) and germline-enriched genes (obtained from Tintori et al ^8^) we found minimal overlap with only 2 out of 245 P-granule transcripts overlapping with the 53 germline-enriched genes (Fisher’s exact test, p = 1.0, Figure S5B and Table S3). The overlap did increase during development, with 31/326 P-granule transcripts overlapping with 160 germline-enriched genes at the four-cell stage (p < 0.001) and 125/389 P-granule transcripts overlapping with 496 germline-enriched genes at the eight-cell stage (p < 0.001; Figure S5B). This progressive enrichment occurs in concert with more transcripts becoming germline-enriched as development proceeds and likely reflects selective degradation these transcripts in somatic lineages ^21,50,58^ rather than active localization to P-granules in the germline.

To further investigate the temporal relationship between P-granule localization and translational repression, we analyzed transcripts that were shown to transition from non-P-granule to P-granule localization during early embryogensis⁵³. While previous studies established that P-granules are not required for translational repression and that poorly translated mRNAs can accumulate in P-granules, these analyses were limited by either focusing on select transcripts^57^, or using pooled embryos across multiple developmental stages^22^, potentially masking stage-specific dynamics. By analyzing 81 transcripts that transition from non-P-granule to P-granule localization^53^ between the 2-cell and 4-cell stages, we observed a significant drop in TE between the 1-cell and 2-cell stages (median difference = −0.4, Wilcoxon rank test, P = 6.28 × 10⁻⁵), before their P-granule localization, with no significant change in TE coinciding with their movement into P-granules (p = 0.6344; Figure S5C and 4C). Thus, for these transcripts, translational repression precedes P-granule localization. A second class of transcripts showed P-granule localization beginning at the 8-cell stage^53^. These 70 transcripts underwent a more gradual decline in TE, but this decline still started before P-granule localization was observed (Figure 4C). We conclude that across the transcriptome, translational repression consistently precedes P-granule localization during early embryogenesis.

Examining P-granule localization across a broader range of developmental time, a previous analysis identified two groups of P-granule transcripts. Group I maintains stable P-granule association throughout primordial germ cell development, while Group II shows transient P-granule localization during early embryogenesis^40^. Group I mRNAs showed consistently lower translational efficiency compared to Group II transcripts across the stages we examined (8-cell medians: −1.34 vs −0.765, p = 2.21e-5, Figure 4D). GO enrichment analysis from a previous study showed that Group I transcripts were enriched for terms related to germline development and reproduction, whereas Group II transcripts were enriched for terms related to metabolic processes^40^, suggesting distinct functional roles for these transcript classes. Together, these findings indicate that translational repression in the germline occurs independently of P-granule association and reveals that P-granules contain functionally distinct transcript populations with different degrees of translational control.

### 2.5. OMA-1 Regulates Translational Efficiency in a Stage-Specific Manner

Our analysis to this point has revealed dynamic changes in TE across developmental stages, which correlate with cell lineage and subcellular mRNA localization. In *C. elegans,* several maternally deposited RNA-binding proteins contribute to spatiotemporally regulated protein expression during early embryonic development^18,27,36,59^. To identify which RNA-binding proteins are most likely to directly regulate translation at the stages we analyzed, we used linear regression analysis (Table S4) to ask how strongly TE correlated with experimental RNA-protein binding data (OMA-1 ^34^ and GLD-1 ^60^) or the presence of binding motifs in transcripts (POS-1 ^61^, MEX-3 ^62^, GLD-1 ^60^ and LIN-41 ^32^) (Figure S6; Methods). OMA-1 binding showed the strongest correlation with translational repression (Figure S6). OMA-1-bound transcripts showed significantly lower TE in the 1-cell stage (median TE: −0.823 vs 0.254; Wilcoxon rank test, p = 8.09e-62) and 2-cell stage (median TE: −0.235 vs 0.177; p = 1.05e-8) (Figure 5A). This repressive effect weakened in later stages, becoming non-significant at the 4-cell stage (median TE: 0.162 vs 0.180; Wilcoxon rank test, p = 0.283) and showing a slight reversal at the 8-cell stage (median TE: 0.293 vs 0.128; p = 0.0171) (Figure 5A). This developmental progression of TE effects correlates with the known degradation of OMA-1 during the 2-cell stage (Figure 5B). Other RNA-binding proteins showed distinct patterns: POS-1, GLD-1, and LIN-41 binding motifs were associated with increased TE across all stages (1-cell stage: coefficients = 0.19, 0.22, and 0.32 respectively, all p < 1e-4), while MEX-3 showed a mild correlation with repression that was only significant at the 2-cell stage (coefficient = −0.10, p = 0.027; Figure S6).

**Figure 5:**
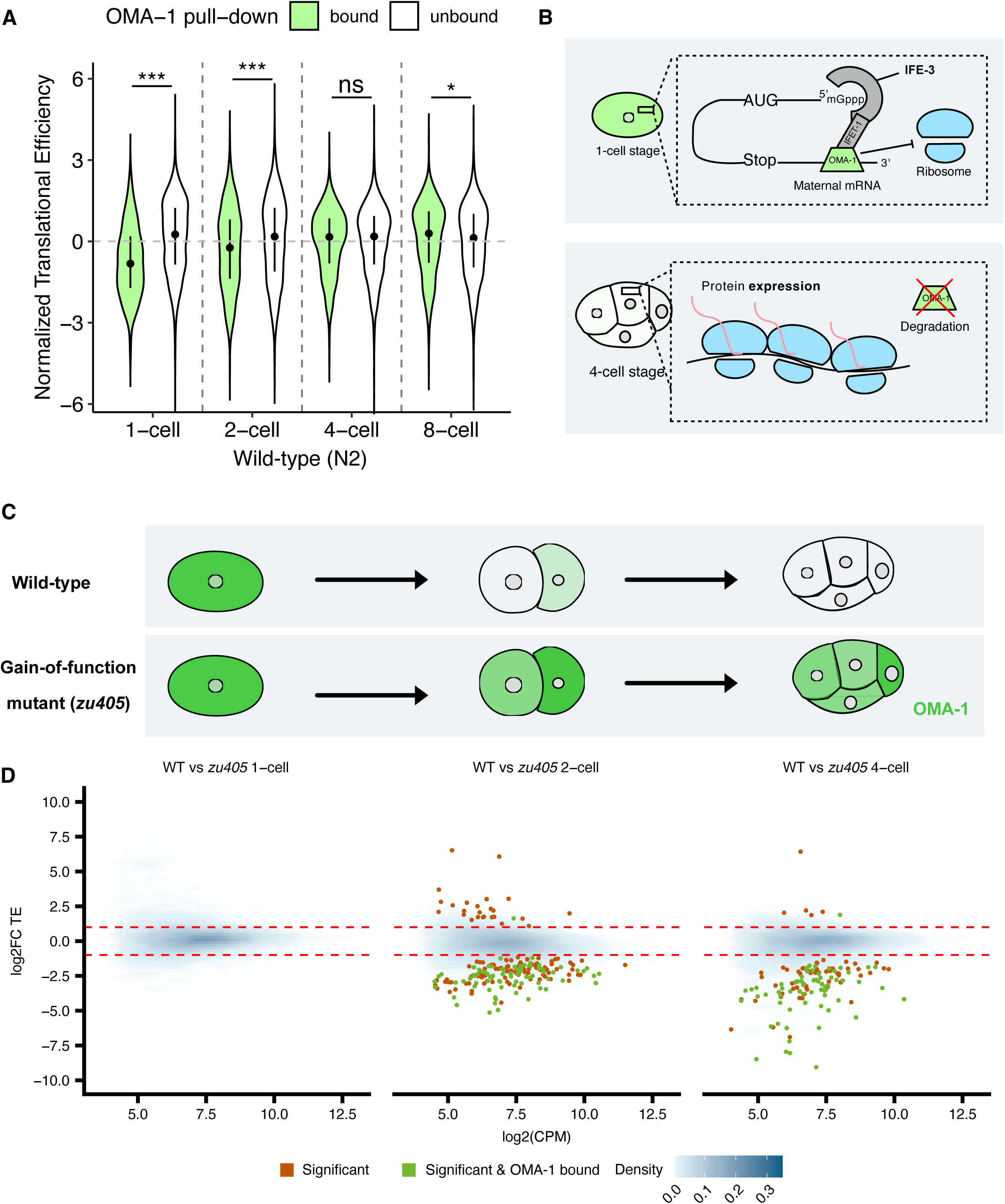
OMA-1 Regulates Translational Efficiency in a Stage-Specific Manner. **(A)** Violin plots in illustrate the distribution of normalized translational efficiency for OMA-1-bound (green) and unbound (white) transcripts (as identified by Spike et al ^34^) across four early *C. elegans* embryonic stages (1-cell, 2-cell, 4-cell, and 8-cell), identified from previous identified from previous OMA-1 associated RNA microarray data. The width of each violin represents the probability density of efficiency values. **(B)** Schematic diagram depicting OMA-1’s regulatory mechanism in early embryogenesis, showing OMA-1 inhibiting translation of maternal transcripts at the 1-cell stage, followed by its degradation by the 4-cell stage, which allows translation of previously repressed transcripts. **(C)** Illustrates the difference in OMA-1 degradation between wild-type and *zu405* mutant embryos. **(D)** A pairwise comparison of translational efficiency between wild-type and mutant at each stage. This is represented as log_2_ fold change in translational efficiency (y-axis) against the mean of normalized counts (x-axis). A blue density plot represents the overall distribution of genes, with intensity corresponding to point density. Orange points indicate transcripts with significant expression changes (FDR < 0.2), while green points highlight known OMA-1-bound transcripts (as identified by Spike et al ^34^) with significant changes in translational efficiency.

Given OMA-1’s striking repressive signature and previous studies showing its role in translationally regulating several specific transcripts during oocyte maturation and early embryogenesis^13,18,34–36,59,63^, we sought to globally characterize OMA-1’s effects on translation during early embryonic development. We carried out Ribo-ITP and RNA-seq experiments using a gain-of-function mutant of *oma-1(zu405)* (Figure S7). This mutant harbors a P240L mutation preventing timely OMA-1 degradation at the two- and four-cell stages and resulting in dominant, temperature-sensitive embryonic lethality^31^ (Figure 5C). We compared TE in *oma-1(zu405)* embryos, raised at the non-permissive temperature, to wild-type embryos (Figure 5D). At the one-cell stage, there were no significant differences (|log_2_FC| >1 and FDR < 0.2), as expected given OMA-1’s presence in both. However, at later stages, we observed widespread aberrant translational repression in the *zu405* mutant, affecting 205 transcripts at the 2-cell stage and 125 at the 4-cell stage (|log_2_FC| >1 and FDR < 0.2; Figure 5D). Interestingly, only a subset of these repressed transcripts (79 at 2-cell and 66 at 4-cell stages) were among the 815 previously identified OMA-1-associated transcripts from adult worms^34^ (green points in Figure 5D). This discrepancy likely reflects both embryo-specific OMA-1 binding not captured in the adult-based pulldown data and indirect regulation through OMA-1’s effects on other RNA regulatory factors. Surprisingly, we also observed relative upregulation (log_2_FC >1 and FDR <0.2) of a small number of transcripts in *oma-1(zu405)* mutants at the 2-cell (26) and 4-cell (6) stages, respectively. These findings highlight the complex and far-reaching effects of OMA-1 on the translational landscape of early *C. elegans* embryos, revealing both direct and potentially cascade effects on gene regulation.

### 2.6. Failure of translational remodeling in oma-1(zu405) reveals diverse regulatory mechanisms

While our initial analysis revealed global TE changes between wild-type and *oma-1(zu405)* embryos at specific stages, we next examined how the temporal progression of translation was altered within the mutant embryos themselves. The *oma-1(zu405)* mutant is 100% embryonic lethal at the non-permissive temperature; embryos fail to specify tissues properly and do not undergo embryonic morphogenesis^31^. By analyzing stage-to-stage transitions in *oma-1(zu405)*, we sought to understand OMA-1’s role in orchestrating the normal developmental program of translational control. Strikingly, there were only 35 genes that changed significantly (|log_2_FC|>1 and FDR <0.2) in TE from the 1-cell to 2-cell stage and only 3 genes from the 2-cell to 4-cell stage in *oma-1(zu405)* (Figure 6A), in contrast to wild-type embryos where hundreds of genes undergo changes in TE between these stages (Figure 2A). In contrast to wild-type embryos, where OMA-1-bound transcripts showed a progressive increase in translational efficiency (TE) over time, in *zu405* mutants the below-average TE for OMA-1-bound transcripts was maintained (Figure S7F, compare to Figure 5A). The median normalized TE of OMA-1 bound transcripts was significantly lower than that of unbound transcripts in 4-cell *oma-1(zu405)* embryos (−0.122 bound vs 0.212 unbound, p = 1.19e-11), in contrast to wild-type where this difference was not significant (0.162 bound vs 0.180 unbound, p = 0.283). Thus, the failure to degrade OMA-1 in the mutant appears to stall the developmental program, preventing the normal translational remodeling that should occur by the 2-cell and 4-cell stages.

**Figure 6:**
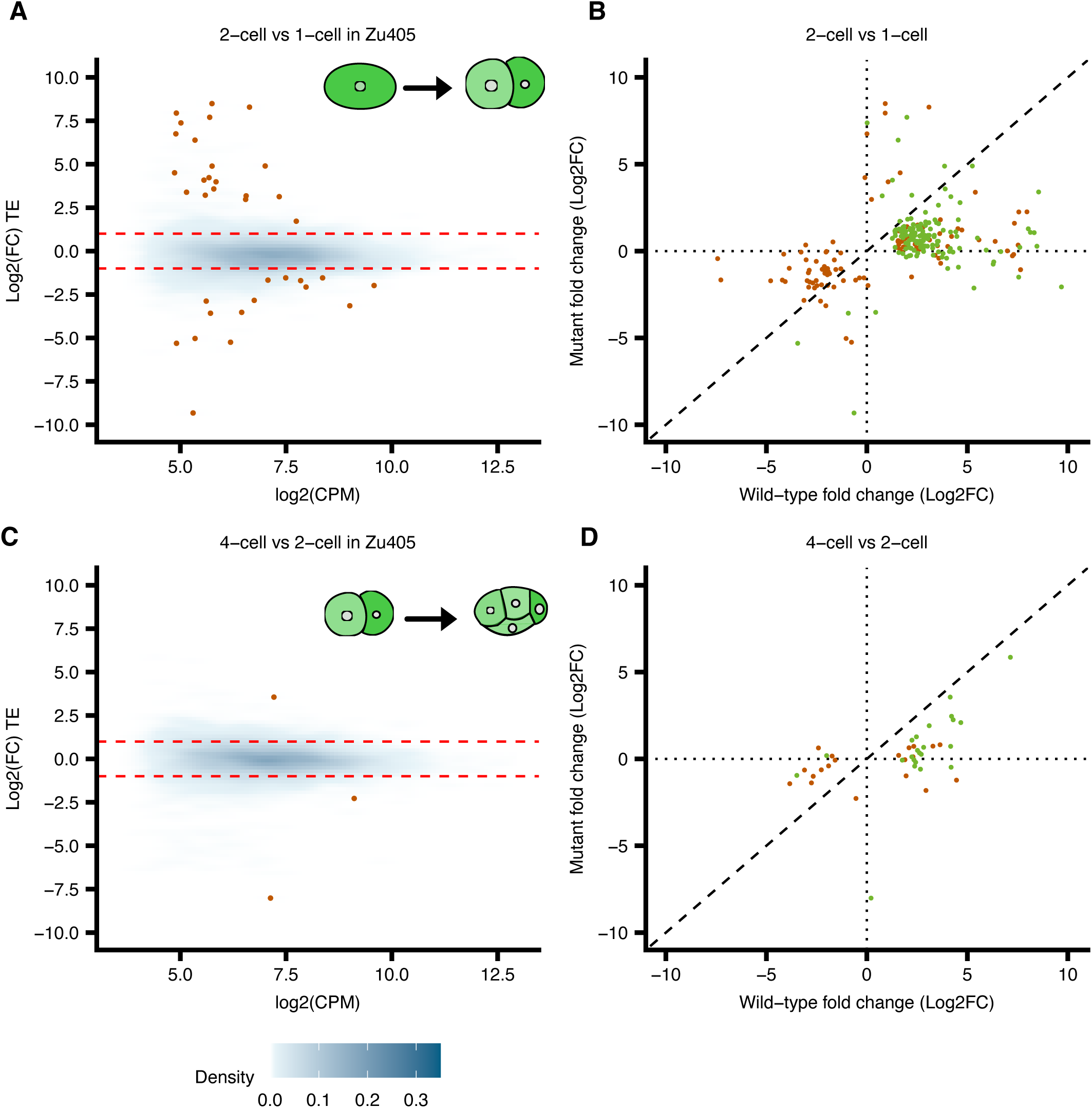
Failure of translational remodeling in oma-1(zu405) reveals diverse regulatory mechanisms. **(A)** MA plot showing changes in TE between 1-cell and 2-cell stages in *zu405* mutant embryos. The log_2_ fold change in TE (y-axis) is plotted against the mean of normalized counts (x-axis). **(B)** Scatterplot comparing log_2_ fold changes in TE between *zu405* mutant (y-axis) and wild-type (x-axis) for genes significantly changed in both genotypes during the 1-cell to 2-cell transition. Points in green indicate known OMA-1 bound transcripts. **(C)** MA plot depicting TE changes between 2-cell and 4-cell stages in *zu405* mutant embryos, with log_2_ fold change in TE (y-axis) plotted against mean normalized counts (x-axis). **(D)** Scatterplot comparing log_2_ fold changes in TE between *zu405* mutant (y-axis) and wild-type (x-axis) for genes significantly changed in both genotypes during the 2-cell to 4-cell transition. Points in green indicate known OMA-1 bound transcripts^34^. In all plots, the blue density overlay represents the overall distribution of genes, with intensity corresponding to the density of data points

To further elucidate which genes might contribute to this developmental stalling, we compared the log fold change in TE between wild-type and mutant embryos for transcripts that showed significant TE changes. Intriguingly, we found that transcripts exhibiting increased TE from the 1-cell to 2-cell stage in wild-type embryos, but no significant change in mutants, were enriched for previously identified OMA-1-bound transcripts (136 out of 179 transcripts; Figure 6B, green points). This pattern suggests that many translationally upregulated transcripts at the 2-cell stage are direct OMA-1 targets that are aberrantly repressed in *zu405*. Conversely, among the transcripts that decreased in TE from 1-cell to 2-cell stage in wild-type embryos but remained unchanged in mutants, none of the downregulated genes was a known OMA-1 binder (Figure 6B). A similar pattern was observed for the 2-cell to 4-cell transition (Figure 6D). We note that changes in mRNA abundance were limited across all conditions, indicating that the majority of these changes in TE arise due to altered ribosome occupancy (Figure S8D and S8E). Additionally, we identified several potential non-OMA-1-bound transcripts with altered TE in mutants (Figure 6A and 6B). These changes likely reflect indirect effects mediated by OMA-1’s regulation of transcripts encoding RNA regulatory factors.

To better understand what types of transcripts are regulated by OMA-1, we moved beyond the gene-by-gene statistical analysis (Figures 5 and 6) to a group-based approach similar to our analysis of wild-type data (Figure 3). This analysis uncovered a spectrum of translational changes among known OMA-1-bound transcript in the mutant condition, revealing four distinct response patterns (Figure 7A). We observed the enrichment of specific biological functions among the detected clusters (Figure 7B). Cluster I, which showed increased TE in the OMA-1 mutant, is enriched for transcripts associated with membrane-bound organelles. Cluster II maintained stable translation regardless of OMA-1 status and was enriched for basic cellular ‘housekeeping’ processes. Cluster III exhibited a minor decreased TE in OMA-1 mutants and was enriched for RNA processing and nuclear functions. Cluster III included *zif-1,* which was shown to be translationally repressed in *zu405* mutants^36^, and our data also observes a similar repression (Figure S8F). Most strikingly, Cluster IV, which showed the strongest translational repression in the mutant, is enriched for developmental processes, particularly tissue development and cell fate commitment, with notable enrichment for Wnt signaling transcripts including *mom-2*, a key Wnt ligand whose translation is dramatically affected in the *oma-1(zu405)* background as confirmed by fluorescent reporters^18^ (Figure S8F). The emergence of these four clusters, each with distinct temporal translational patterns and enriched biological processes, highlights the complex and varied roles of OMA-1 in regulating its target transcripts, which might be explained by OMA-1’s known interactions with several other RNA-binding proteins^18,32^. Furthermore, the detection of OMA-1-bound transcripts whose translation is unaffected in the mutant suggests that these transcripts may only be regulated at other stages (e.g. oogenesis), or that OMA-1 binding alone is insufficient to dictate their translational regulation during early embryogenesis. Together, these results reveal that OMA-1 differentially regulates distinct functional groups of transcripts, with particularly strong effects on genes involved in development and cell fate decisions, while having more modest or indirect effects on basic cellular processes.

**Figure 7:**
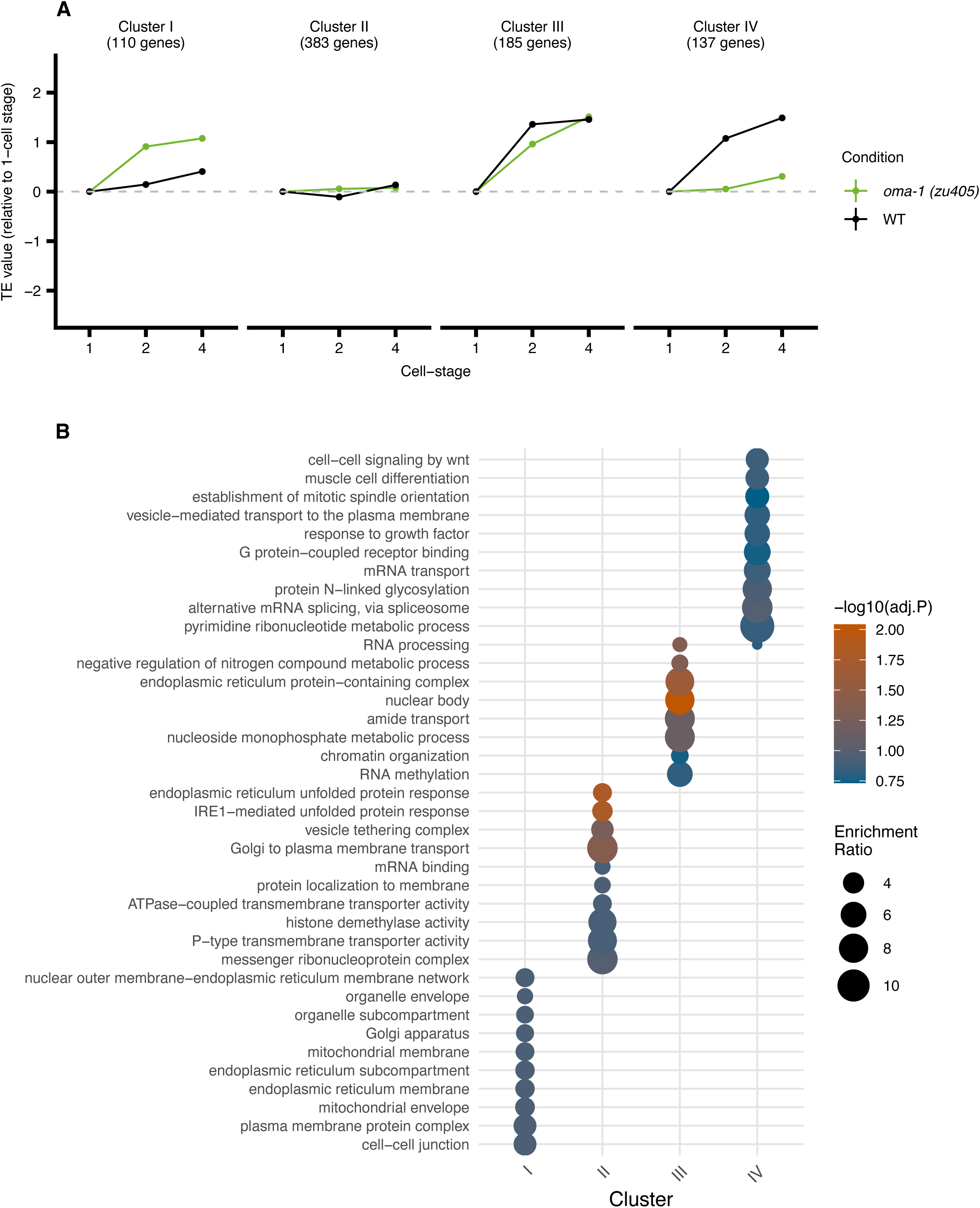
Temporal clustering reveals diverse OMA-1-mediated translational regulation. **(A)** Line plot showing four clusters of OMA-1-bound transcripts identified based on changes in TE. The y-axis represents TE values normalized to the 1-cell stage, and the x-axis shows developmental stages. Each line represents the mean TE trajectory for a cluster in both wild-type (black) and mutant embryos (green). Error bars represent the standard error of TE values for all genes within each cluster**. (B)** Dot plot visualizing Gene Ontology (GO) enrichment analysis for the four identified clusters. The y-axis displays enriched GO terms, while the x-axis shows the clusters. Dot size corresponds to the enrichment ratio and color intensity indicates the adjusted p-value of enrichment.

## 3. Discussion

Translational regulation allows precise control of protein expression in the absence of active transcription in early embryonic development across various species. Our study provides the first comprehensive, high-resolution view of translational dynamics during the initial cell divisions in *C. elegans* embryos, revealing extensive regulation of maternal mRNAs. By leveraging ribosome profiling via isotachophoresis (Ribo-ITP)^23^, we overcame previous technical limitations in analyzing low-input samples from precisely staged embryos. This approach enabled us to generate high-quality ribosome footprint data from individual embryonic stages with unprecedented temporal resolution. Combined with low-input RNA-seq, we uncovered global patterns of translational control that shed light on how cell fates are specified prior to zygotic genome activation. Our findings not only extend our understanding of early embryogenesis in *C. elegans* but also provide insights into the fundamental mechanisms of post-transcriptional regulation in early animal development.

Previous studies were limited to RNA-seq and mass spectrometry analyses of oocytes, 1-cell, and 2-cell stages, and did not directly investigate translational regulation^49^. Our study demonstrates that ribosome recruitment onto mRNA is poorly correlated with mRNA abundance in the *C. elegans* early embryos. The correlation between ribosome occupancy and mRNA levels varies across species in a way that appears to correlate with the rate of embryonic development. *Drosophila* embryos (Pearson correlation = 0.348)^64^ develop rapidly, with the entire embryogenesis completed in about 24 hours, while mouse embryogenesis (Spearman correlation ranging from 0.7 to 0.8)^23^ takes approximately 20 days. This spectrum of developmental rates and strategies extends to vertebrates as well. Zebrafish ^65^ and *Xenopus* ^66^ develop faster than mammals, completing embryogenesis in 3-4 days with reduced reliance on zygotic transcription^67–69^. We suspect that, like *C. elegans*, these faster-developing vertebrates will depend more heavily on maternal factors and post-transcriptional regulation during early embryogenesis.

The relationship between developmental tempo and translational regulation is further illustrated by comparing *C. elegans* with the nematode *Ascaris suum*. Despite similarities in early embryonic cellular organization, *A. suum* initiates zygotic genome activation at the 1-cell stage, prior to pronuclear meeting^70^. Surprisingly, many of the maternal transcripts required in *C. elegans* are zygotically produced in *A. suum*. This difference may be attributed to the vastly different cell cycle durations: approximately 24 hours in *A. suum* (similar to mouse) compared to 10-15 minutes in *C. elegans*. These observations suggest that organisms undergoing rapid cell division may co-opt translational regulation to meet changing protein demands more efficiently, with *C. elegans* representing an extreme case of rapid development and reliance on post-transcriptional regulation among these model organisms.

During early embryonic development, maternal mRNAs are often localized to specific regions of the egg or embryo, establishing developmental axes and patterns. These localized mRNAs, when translated, produce proteins that initiate and maintain developmental gradients^71,72^ and specify cell fates^10,73^. For instance, in many organisms, mRNAs localized to the animal and vegetal poles of the egg are responsible for determining the anterior-posterior and dorsal-ventral axes^72^. Additionally, some localized mRNAs are crucial for germ plasm assembly and germ cell development^74^. By comparing TE values for mRNAs with known subcellular localization in *C. elegans*^57^, we detected correlations between the localization of transcripts and their respective TE. This relationship was particularly apparent for mRNAs localized to specific cellular compartments, such as the cell periphery and in germline-specific ribonucleoprotein complexes. Our analyses revealed that mRNAs localized to the periphery were relatively translationally active, while those in P granules were translationally repressed. Interestingly, prior studies suggested that the localization of some mRNAs to the cell periphery may require partial translation to occur^19,57^. These findings suggest that subcellular positioning of transcripts may serve as a key mechanism for fine-tuning gene expression in diverse cellular processes and developmental stages across various species.

Our analysis further builds on recent insights^22,40,57^ on the temporal relationship between translational repression and P-granule association in the germline lineage. While germline-localized transcripts showed consistently lower translational efficiency compared to somatic transcripts, this repression preceded their enrichment in P-granules. The lack of P-granule transcript enrichment at the two-cell stage, despite strong translational repression, coupled with the observation that transcripts transitioning to P-granule localization were already translationally repressed, suggests that initial translational control in the germline may operate prior to P-granule association. These temporal observations implicate a model where repression is established before transcripts become enriched in P-granules. Future studies using techniques such as the SunTag ^75,76^ could provide direct visualization of translation events in relation to P-granule localization to further validate this model.

P-granule transcripts can be classified into two groups based on their localization dynamics. Group I transcripts maintain stable P-granule association through primordial germ cell development, while Group II transcripts show transient association during early embryogenesis^40^. The stronger translational repression observed in Group I compared to Group II transcripts suggests that P-granules may serve as platforms for maintaining or reinforcing established translational programs. This two-step model, where initial translational repression might occur in the cytoplasm and is followed by P-granule-mediated maintenance for Group I transcripts, provides new insights into how germline identity is established and maintained during early development. The role of P-granules in maintaining translational repression of Group II transcripts, however, remains to be determined.

Intriguingly, we observed that OMA-1-bound transcripts ^34^ underwent varying levels of change in TE in the OMA-1 gain-of-function mutant. If these transcripts were solely controlled by OMA-1, we would expect uniform changes across all targets. Instead, we observe context-dependent variations, likely influenced by the cell stage and possibly by other RNA-binding proteins working in concert with OMA-1^13,32,33^. Most surprisingly, while OMA-1 predominantly functions as a translational repressor, a subset of transcripts showed increased TE in the gain-of-function mutant (Figure 5D, orange dots above the line), suggesting OMA-1 can activate translation of some transcripts. This activity could occur through direct activation of specific targets, as previously demonstrated for SPN-4 and MEG-1^32^, or through an indirect mechanism where OMA-1 inhibits other translational repressors. Additionally, it is noteworthy that only a minority of OMA-1-bound transcripts (∼10%) exhibit strong translational repression at the stages we analyzed, further supporting the hypothesis that effective OMA-1-mediated regulation requires cooperative action with stage-specific cofactors. The clustering analysis in Figure 7A illustrates these diverse regulatory behaviors. These findings emphasize the need to consider both direct and indirect regulatory mechanisms and the combinatorial effects of RNA-binding proteins in early embryogenesis.

Altogether, our work has uncovered modes of translational regulation that shape protein expression during early embryogenesis. Our high-resolution temporal analysis of translational dynamics provides insight into how post-transcriptional regulation directs cell fate decisions. This technical advance enables systematic identification of global regulatory roles for RNA binding proteins during development. The *C. elegans* model system’s unique amenability to both genetic manipulation and global analyses makes it ideally suited for these studies, allowing us to combine our novel approaches with classical genetics in ways not possible in other organisms. This comprehensive dataset will serve as a valuable resource for the *C. elegans* community, enabling future studies of translational control in early development and cellular differentiation.

### 3.1. Limitations of the Study

First, we used ribosome occupancy normalized to mRNA abundance as a proxy for translational efficiency, but note that our measurements cannot distinguish between actively translating ribosomes and stalled ribosomes on transcripts. Additionally, our approach lacks single-molecule resolution, meaning we cannot distinguish whether changes in ribosome footprint abundance represent uniform translation across all copies of a particular mRNA, or heterogeneous translation where some copies are heavily translated while others remain untranslated. This distinction may be particularly relevant in early embryos where spatial organization of mRNAs plays important developmental roles. Second, the compositional nature of any sequencing-based measurement provides only relative estimates rather than absolute measurements, which should be carefully considered when interpreting the results. Third, our RNA sequencing approach relied on poly-A selection, thereby excluding non-polyadenylated transcripts such as histones from our analysis. Fourth, our current experimental design, which examined whole embryos at the 1-cell, 2-cell, 4-cell, and 8-cell stages, lacks spatial resolution at the lineage level. The calculated translation efficiency values represent averages across different cell lineages rather than lineage-specific measurements. Future experiments could address this limitation by investigating translational regulation within specific lineages of the *C. elegans* embryo to better understand lineage-specific translational control.

## 4. Acknowledgments

This work was supported by National Institutes of Health grants [HD110096, GM150667] and a Welch Foundation grant [F-2027-20230405] (C.C.); and by NIH GM138443 (D.D.). We thank N. Stolpner for contributions to early project conceptualization, D. Park for experimental troubleshooting assistance, R. Tachibana for analysis validation, I. Hoskins for computational support with 3’UTR analysis, M. Modi for valuable discussions on statistical approaches and data visualization, and Sophia Tintori and Bob Goldstein for critical reading of the manuscript. We thank Germano Cecere for sharing their translational efficiency data. We also thank members of the Cenik and Dickinson labs for helpful discussions.

## 5. Author contributions

Y.S., D.J.D., and C.C. conceptualized the study. Y.S. and V.G. performed ribosome profiling experiments and data analysis with guidance from C.C.. Y.S. analyzed P-granule localization data. C.H. performed imaging experiments and analysis under supervision from D.J.D.. Y.S. performed bioinformatics analysis with input from C.C.. Y.S. wrote the manuscript with input from all authors. Y.S. prepared the figures with suggestions from V.G., D.J.D., and C.C.. D.J.D. and C.C. supervised the project.

## 7. Declaration of generative AI and AI-assisted technologies

During the preparation of this work, we used Claude (Anthropic) for three purposes: (1) to improve grammar and clarity of scientific writing, (2) to generate code for data visualization and statistical analysis, which was subsequently reviewed and modified. All AI-generated content was carefully reviewed, edited, and validated by the authors. The authors take full responsibility for the accuracy and completeness of the published manuscript.

## 8. Accession number

The accession number for all libraries reported in this paper is NCBI GEO: GSE281412. Original image data will be deposited in the Texas Data Repository (https://dataverse.tdl.org), with DOI: https://doi.org/10.18738/T8/EFJNGZ.

## 9. Code availability

The code used in the study is available at https://github.com/yss322/Low-input-ribosome-profiling-of-Celegans-embryogenesis

## 10. Supplemental information titles and legends

**Document S1:** Figures S1–S7 and Table S1-S5 description

**Table S1:** Count Data for RNA Sequencing and Ribosome Profiling

**Table S2:** GO annotations Gene Ontology Enrichment Analysis Results for WT

**Table S3:** Fisher’s Exact Test Analysis of P-Granule and Germline-Enriched Gene Overlap

**Table S4:** Linear Regression Analysis of RNA-Binding Protein Effects on Translation Efficiency

**Table S5:** Gene Ontology Enrichment Analysis of OMA-1-Regulated Transcript Clusters

## 11. Supplementary Figures

**Figure S1:**
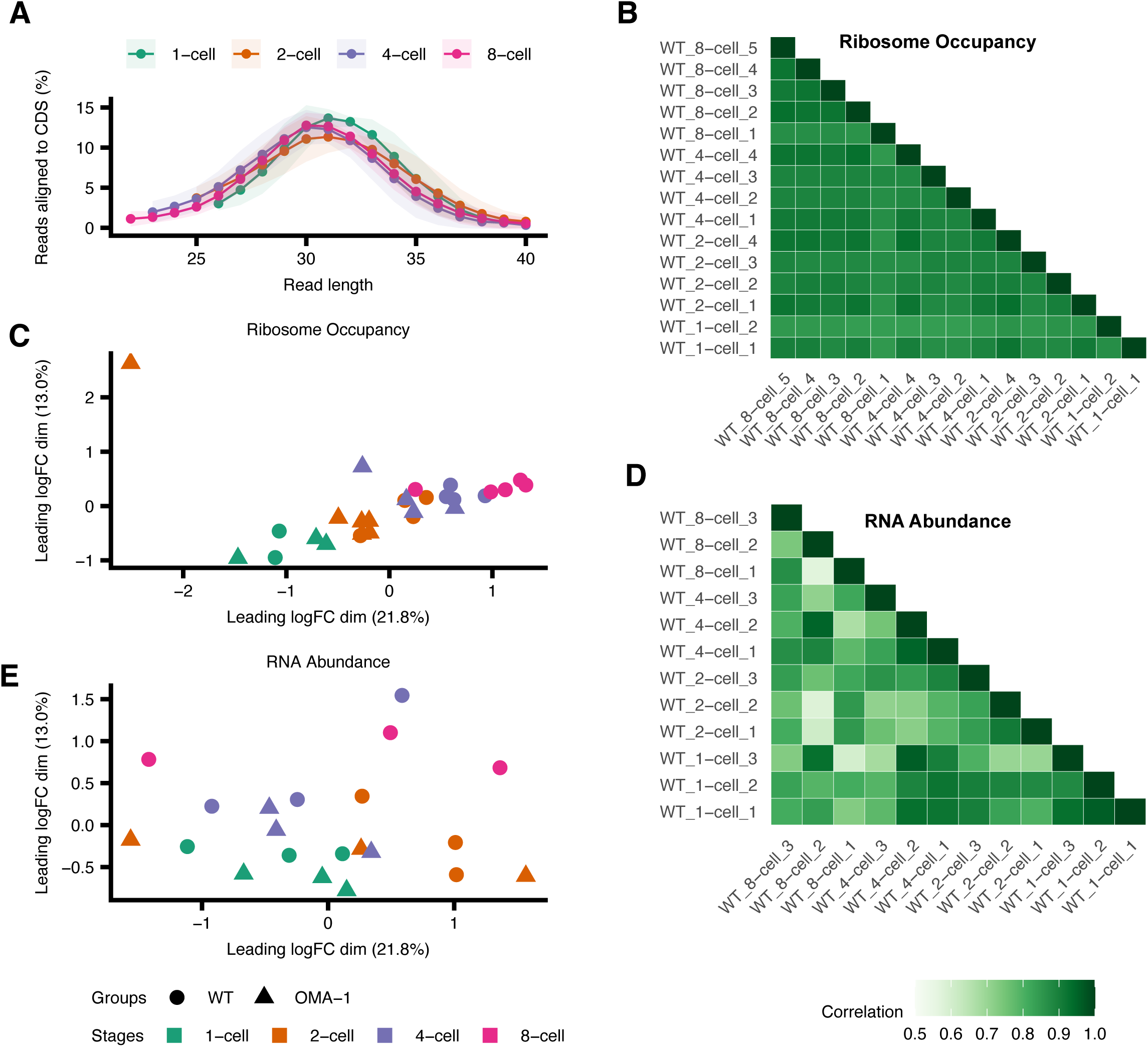
**(A)** Distribution of ribosome profiling read lengths across developmental stages, with shaded areas indicating standard deviation between biological replicates**. (B)** Correlation matrix of ribosome occupancy, where darker colors indicate stronger correlations. **(C)** Multidimensional scaling analysis of Ribo-seq. Axes represent arbitrary units. **(D)** Correlation matrix of RNA abundance between replicates, where darker colors indicate stronger correlations. **(E)** RNA-seq replicates, with shapes denoting WT versus OMA-1 mutants and colors indicating cell stages. Axes represent arbitrary units.

**Figure S2:**
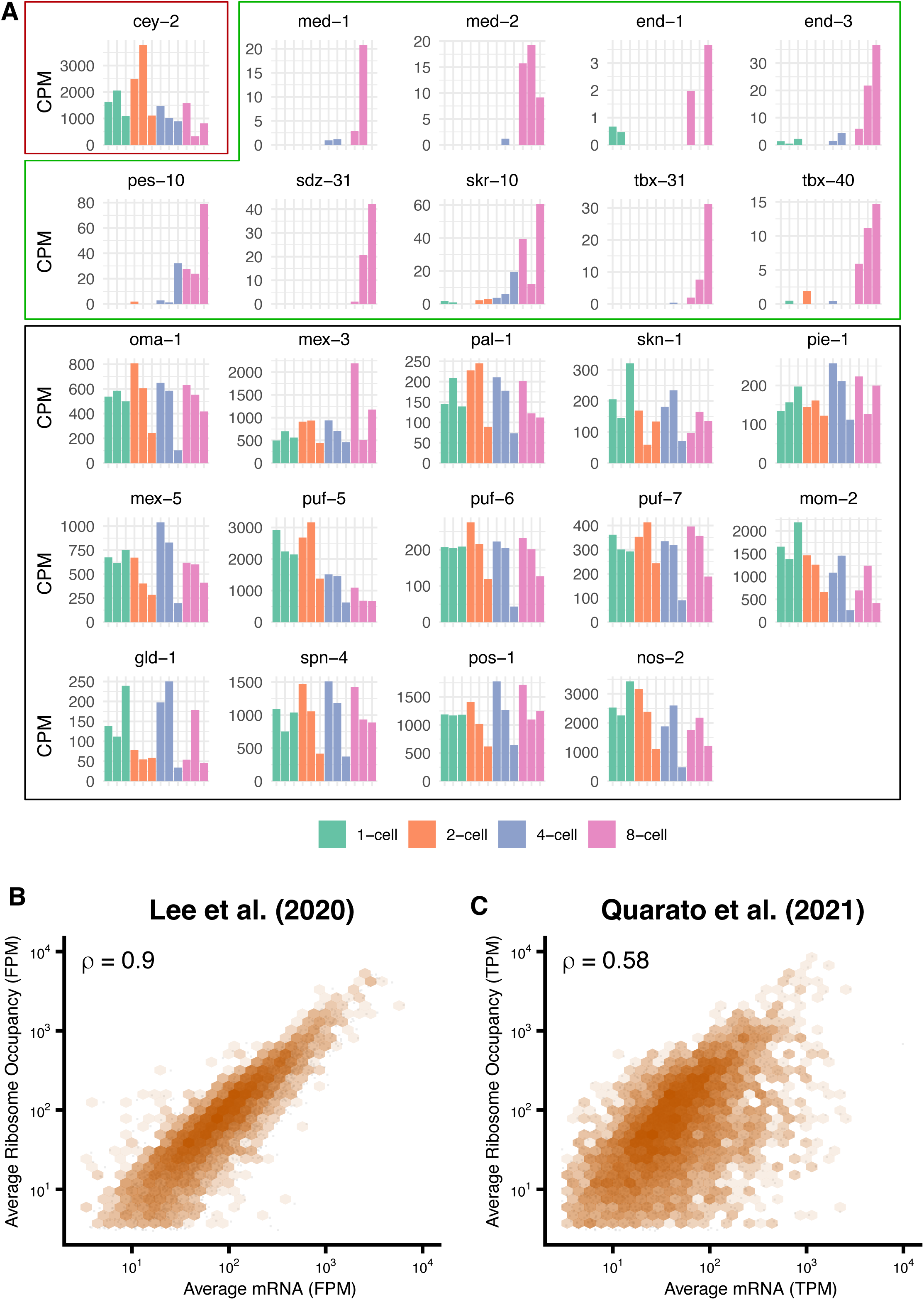
(A) Expression patterns from RNA-seq of marker genes, highlighting cey-2 in red box showing progressive degradation throughout embryogenesis, while genes predominantly expressed at the 8-cell stage are in green box. Other previously identified maternally deposited mRNA are shown in the black box. Each bar in the plot of the same color represents a replicate. Relationship between mRNA abundance and ribosome occupancy from (B) Lee et al. (C) Quarato et al. (2021). The plot shows average mRNA levels versus average ribosome occupancy on log10 scales. Gray dots represent individual genes with hexagonal binning (orange) indicating data density. The Spearman correlation coefficient (rho) quantifies the relationship strength

**Figure S3:**
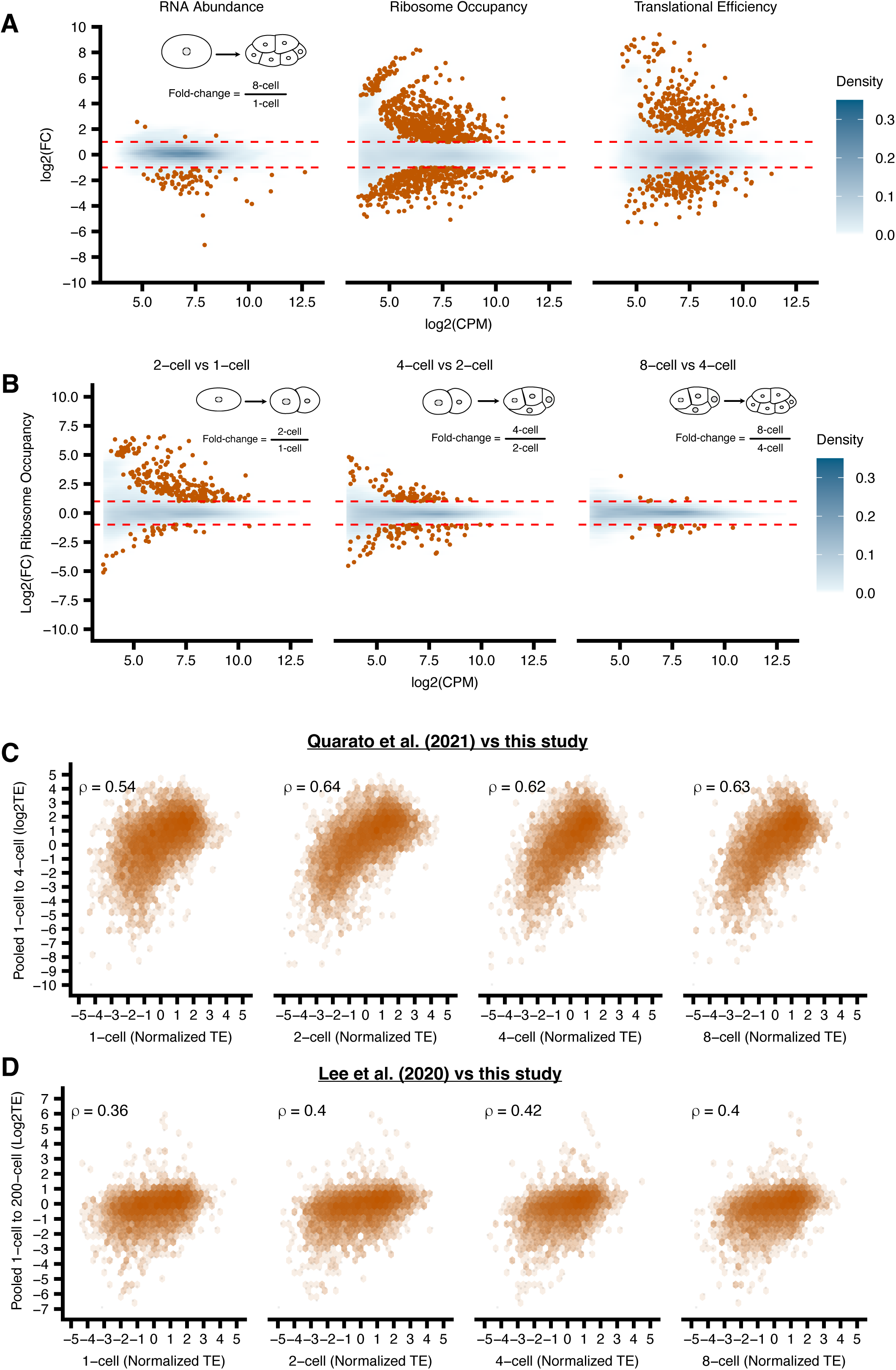
**(A)** Mean difference plots comparing RNA abundance, ribosome occupancy, and translational efficiency between 8-cell and 1-cell stage wild-type embryos. X-axis shows expression level (log2CPM), Y-axis shows log_2_ fold change. Mean difference plots comparing gene expression between sequential *C. elegans* embryonic stages **(B)**1-cell to 2-cell, 2-cell to 4-cell, and 4-cell to 8-cell. The log2 fold change in ribosome occupancy (y-axis) is plotted against the mean of normalized counts (x-axis). Blue density plot represents the overall distribution of genes with the intensity corresponds to the density of points. The orange points indicate transcripts with significant expression changes (FDR < 0.2 and log_2_FC >1 or < −1). Blue density indicates concentration of data points. Orange points highlight significant changes (|log_2_FC| > 1, FDR < 0.05). Red dashed lines indicate ±1 log_2_FC thresholds. Correlations between stage-specific and pooled embryonic translational efficiencies. **(C)** Comparison of normalized TE at the same developmental stages with pooled 1-cell to 4-cell TE data from Quarato et al. (2021) (y-axes). **(D)** Comparison of normalized translational efficiency (TE) at 1-cell, 2-cell, 4-cell, and 8-cell stages (x-axes) with pooled 1-cell to 200-cell TE data from Lee et al. (2020) (y-axes). Orange hexagonal binning shows data density with Spearman correlation coefficients (rho) indicating relationship strength.

**Figure S4:**
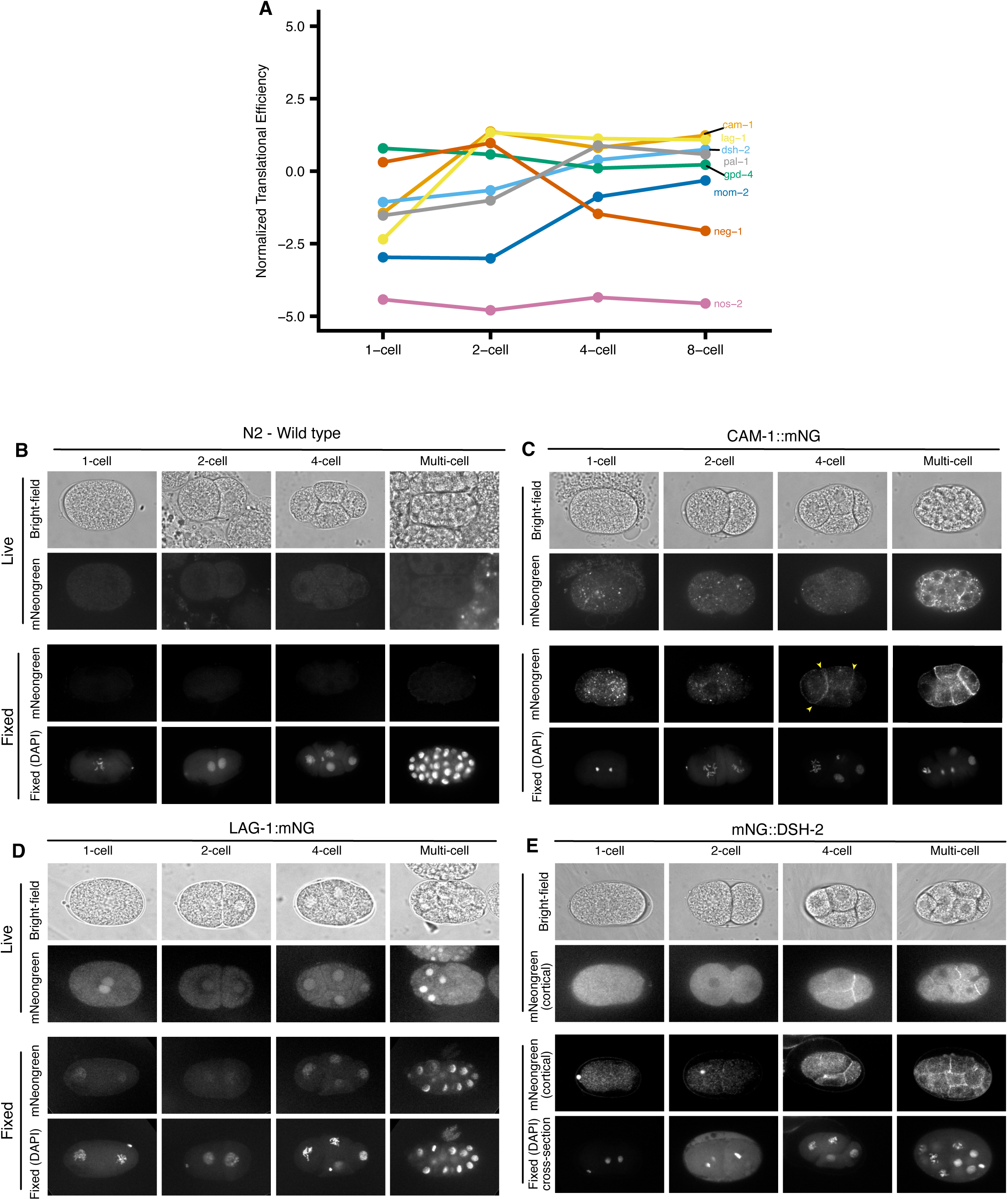
**(A)** Normalized TE profiles of previously identified (PAL-1, MOM-2, NEG-1, NOS-2) and newly identified (CAM-1, LAG-1 and DSH-1) translationally regulated transcripts compared to the housekeeping gene GPD-4. **(B-E)** Localization of newly identified proteins during early embryonic development. **(B)** N2 control, **(C)** CAM-1::mNG, **(D)** LAG-1::mNG, and **(E)** DSH-1::mNG embryos imaged at 1-cell, 2-cell, 4-cell, and multicellular stages. For each genotype: brightfield (top row), live mNG fluorescence (middle row), and fixed samples showing mNG signal with DAPI counterstain (bottom row). Arrowheads indicate cell-cell contact localization of fluorescently tagged proteins in CAM-1::mNG.

**Figure S5:**
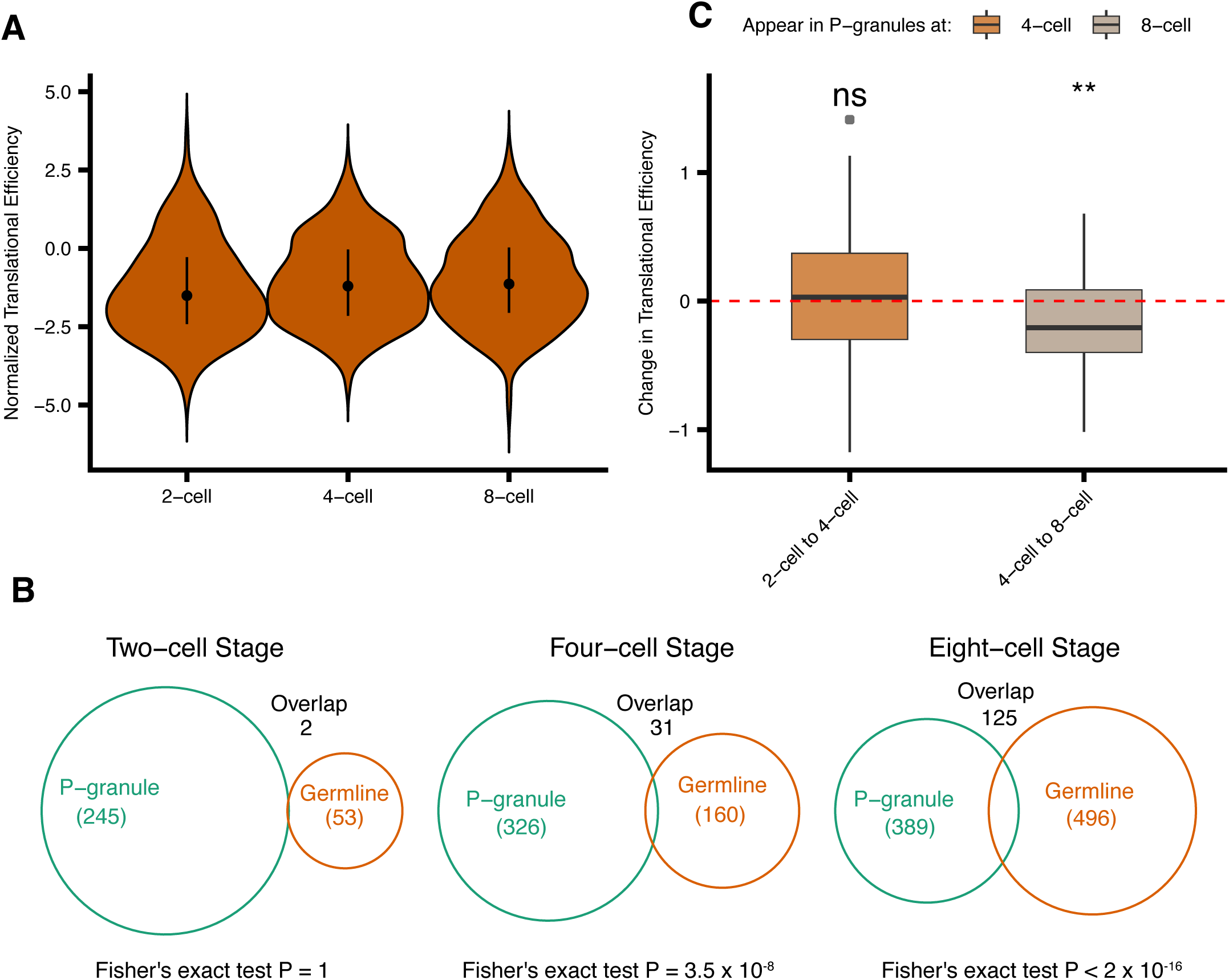
**(A)** Translational efficiency of P-granule localized transcripts across developmental stages. Violin plots show distribution of TE values for P-granule transcripts at 2-cell (n=245), 4-cell (n=326), and 8-cell (n=389) stages, with median and quartiles indicated. **(B)** The overlap between P-granule-associated genes and genes expressed at specific developmental stages in *C. elegans* early embryogenesis. The Venn diagrams illustrate three developmental timepoints: two-cell, four-cell, and eight-cell stage. **(C)** Changes in translational efficiency of transcripts as they transition into P-granules. Box plots show TE changes for transcripts first appearing in P-granules at 4-cell (orange, n=81) and 8-cell stages (brown, n=43). Red dashed line indicates no change in TE. Asterisks denote significant differences from zero (***p < 0.001, Wilcoxon test).

**Figure S6:**
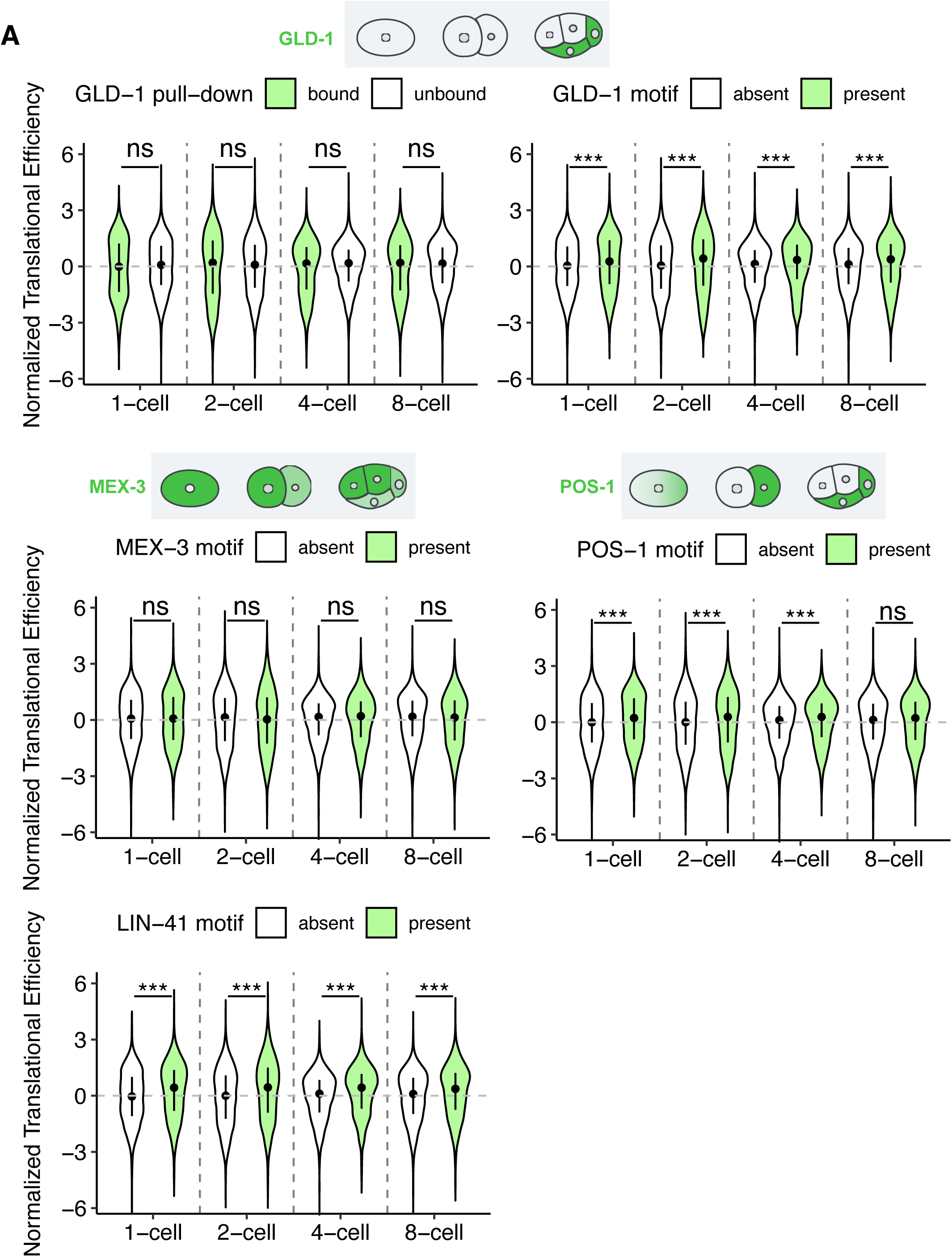
Violin plots showing the distribution of normalized translational efficiency for transcripts across early *C. elegans* embryonic stages (1-cell, 2-cell, 4-cell, and 8-cell). Comparisons are shown between: **(A)** GLD-1-bound versus unbound transcripts from RNA-immunoprecipitation data, and transcripts with present versus absent motifs for **(B)** POS-1, **(C)** MEX-3, **(D)** GLD-1, and **(E)** LIN-41. Green violins represent bound/motif-present transcripts, while white violins indicate unbound/motif-absent transcripts. Plot width indicates probability density of translational efficiency values(***p < 0.001, Wilcoxon test). Illustrations indicate known localization of proteins during early embryogenesis

**Figure S7:**
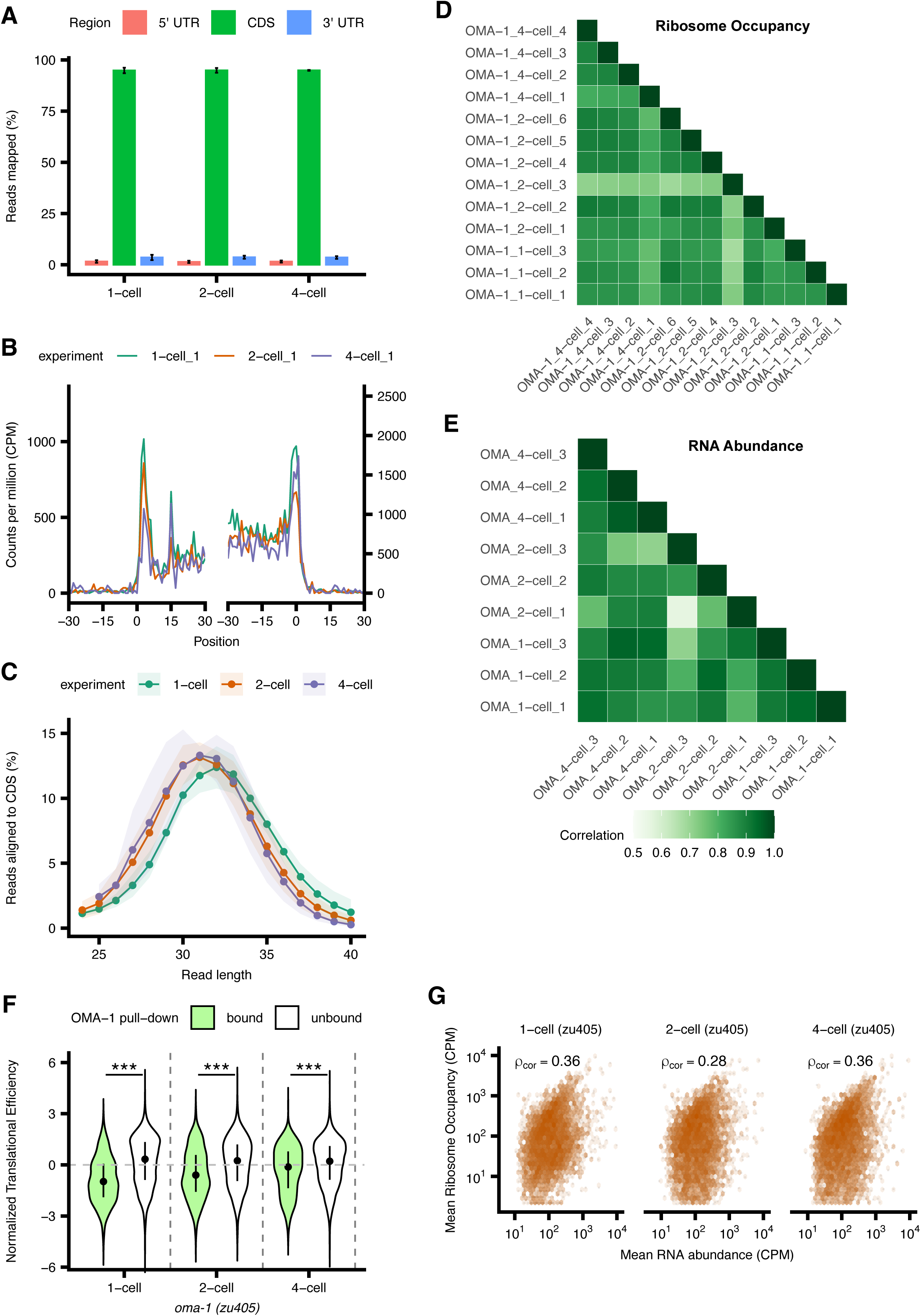
**(A)**The allocation of ribosome profiling reads across genomic features (CDS, 5’ UTR, 3’ UTR) is presented for each stage in *zu405* mutants. **(B)** Ribosome occupancy around the translation start and stop sites in a representative 1-cell, 2-cell and 4-cell *zu405* staged embryo. Translation start (or stop) sites are denoted by the position 0. Aggregated read counts (*y* axis) relative to the start (or stop) sites are plotted after A-site correction **(C)** Distribution of ribosome profiling read lengths across developmental stages, with shaded areas indicating standard deviation between biological replicates**. (D)** Correlation matrix of ribosome occupancy and **(E)** RNA abundance between replicates, where darker colors indicate stronger correlations. **(F)** Violin plots in illustrate the distribution of normalized translational efficiency for OMA-1-bound (green) and unbound (white) transcripts across three early *zu405* embryonic stages (1-cell, 2-cell and 4-cell) identified from previous RNA-immunoprecipitation data. The width of each violin represents the probability density of efficiency values(***p < 0.001, Wilcoxon test). **(G)** Pairwise correlation between ribosome occupancy and RNA abundance at the three stages of early *zu405* embryo development is presented. The mean cpm of ribosome occupancy is plot against the mean cpm of RNA abundance at each stage. Rho (cor) is the corrected spearman correlation based on the reliability r (RNA) and r (Ribo) which are the replicate-to-replicate correlation (methods).

**Figure S8:**
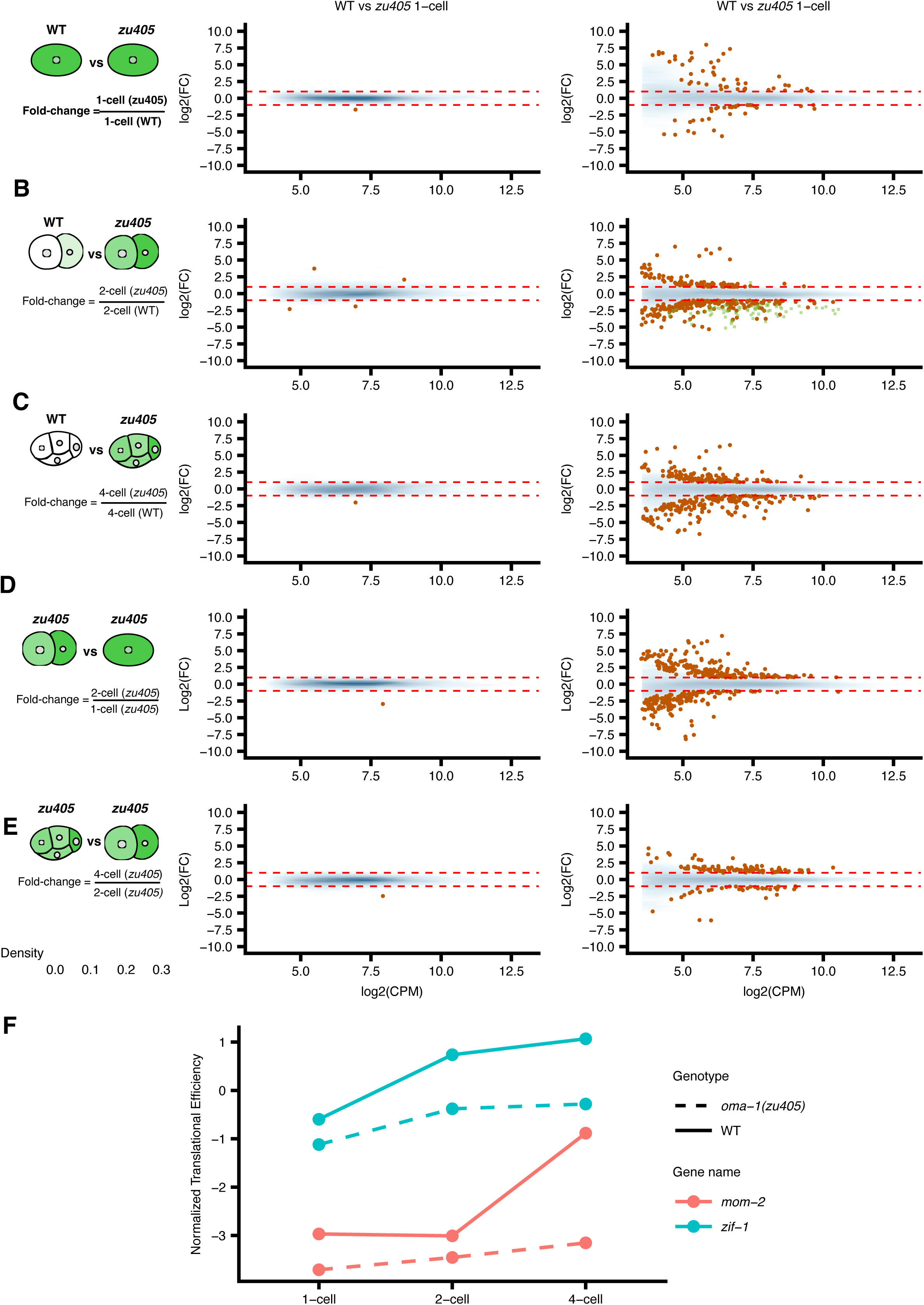
Mean difference plots comparing gene expression between *zu405* mutant and wild-type embryos. **(A-C)** Comparisons between *zu405* and wild-type at 1-cell, 2-cell, and 4-cell stages, respectively. **(D-E)** Stage-to-stage comparisons within zu405 embryos from 1-cell to 2-cell and 2-cell to 4-cell. For all plots, log_2_ fold changes in RNA abundance and translational efficiency (y-axis) are plotted against mean normalized counts (x-axis). Blue density shading indicates the distribution of all genes, with darker shading representing higher point density. Orange points highlight significantly differentially expressed genes (FDR < 0.2 and |log_2_FC| > 1). **(F)** Developmental dynamics of translational efficiency for mom-2 and zif-1 in wild-type and oma-1 mutant embryos. The plot compares normalized translational efficiency (TE) of mom-2 and zif-1 genes across early embryonic stages (1-cell, 2-cell, and 4-cell) in both wild-type (WT, solid lines) and oma-1(zu405) mutant (dashed lines) backgrounds.

## 13. STAR Methods

### Key resources table

| REAGENT or RESOURCE | SOURCE | IDENTIFIER |
| --- | --- | --- |
| Chemicals, peptides, and recombinant proteins |  |  |
| Cycloheximide | Sigma | Cat#C4859-1ML |
| T4 Polynucleotide Kinase | New England Biolabs | Cat#M0201L |
| Sodium Hypochlorite | Sigma | Catalog Number: 239305 |
| Inulin | Thermo | Catalog Number: A18425.18 |
| PVP Powder | Thermo | Catalog Number: 227545000 |
| BME Vitamins | Sigma | Catalog Number: B6891 |
| Chemically Defined Lipid Concentrate | Gibco | Catalog Number: 11905031 |
| Penicillin-Streptomycin | Sigma | Catalog Number: P4333 |
| Schneider's Drosophila Medium | Gibco | Catalog Number: 21720024 |
| Fetal Bovine Serum (FBS) | Gibco | Catalog Number: A3840001 |
| Critical commercial assays |  |  |
| D-Plex Small RNA-seq Kit | Diagenode | Cat#C05030001 |
| NEBNext Single Cell/Low Input RNA Library Prep Kit for Illumina | New England Biolabs | Cat#E6420S/L |
| Deposited data |  |  |
| Raw and analyzed data | This Paper | GEO: GSE281412 |
| *C. elegans* reference files | Liu et al<sup>44</sup> | https://github.com/RiboBase/reference_caenorhabditis-elegans |
| Study code and analysis | This Paper | https://github.com/yss322/Low-input-ribosome-profiling-of-Celegans-embryogenesis |
| Raw image data | This Paper | Texas Data Repository<br>https://doi.org/10.18738/T8/EFJNGZ |
| Experimental models: Organisms/strains |  |  |
| *C. elegans*: Wild type N2 | Caenorhabditis Genetics Center | N2 |
| *C. elegans* OMA-1 mutant strain | Lin R. et al<sup>31</sup> | TX20<br>Genotype: *oma-1(zu405)* IV. |
| *C. elegans* CAM-1::mNG::3xFLAG strain | Heppert et al<sup>56</sup> | LP530<br>Genotype: *cam-1(cp243[cam-1::mNG-C1^3xFlag])* II. |
| *C. elegans* DSH-2::mNG::3xFLAG strain | Heppert et al<sup>56</sup> | LP228 dsh-2(cp51[mNeonGreen::dsh-2]) II |
| *C. elegans* LAG-1::mNG | This study | UTX259 lag-1(dev208([lag-1::mNeonGreen]) IV |
| Software and algorithms |  |  |
| RiboFlow | Ozadam et al<sup>77</sup> | https://github.com/ribosomeprofiling/riboflow |
| UMI-tools | Smith et al<sup>78</sup> | https://umi-tools.readthedocs.io/en/latest/reference/extract.html |
| Bowtie2 | Langmead and Salzberg<sup>79</sup> | http://bowtie-bio.sourceforge.net/bowtie2/index.shtml |
| Cutadapt | Marcel Martin<sup>80</sup> | https://cutadapt.readthedocs.io/en/stable/# |
| ComBat-seq | Zhang et al<sup>81</sup> | https://github.com/zhangyuqing/ComBat-seq |
| edgeR | Robinson et al<sup>82</sup> | https://bioconductor.org/packages/release/bioc/html/edgeR.html |
| clusterProfiler | Yu et al<sup>83</sup> | https://guangchuangyu.github.io/software/clusterProfiler/ |
| Micro-Manager | Edelstein et al<sup>84</sup> | https://micro-manager.org/ |

### 13.1. Experimental model details

#### *C. elegans* maintenance

*C. elegans* strains were maintained on standard NGM plates and fed OP50 E. coli. The N2 (wild type) and LP530 strains were maintained at 20°C and the TX20 strain was maintained at 15°C.

### 13.2. Method Details

#### Embryo collection

For wild-type strains, embryos were collected as follows: Microscope slides were prepared by placing five Eppendorf tube stickers in a row and applying Sigmacote to the surrounding area. After drying under a fume hood for 5-10 minutes, the slides were washed with water and the stickers removed, creating wells. For embryo collection, approximately 20 worms were placed in 100 µL of egg buffer and dissected. 10 µL of bleach was added, and the solution was pipetted for 10 seconds. Eggs were selected and washed five times through 100 µL of SGM in the wells created on the treated slides (taking ∼ 2 mins). After reaching the desired stage, eggs were washed five times in water, then in water with 100 μg/ml cycloheximide (CHX). During the final wash, embryos were inspected using a microscope, and those not at the correct stage were segregated. The correctly staged embryos were transferred to a PCR tube, flash-frozen in liquid nitrogen, and then kept on a metal rack placed on dry ice. More embryos were added to the same PCR tube on dry ice, with the tube being flash-frozen in liquid nitrogen each time new embryos were added. This process was repeated until 9 embryos were collected in a single PCR tube for each stage.

For TX20 *C. elegans* strains, L4/young adult worms were incubated overnight at 20°C prior to collection. Following this incubation, the same collection process as described for wild-type strains was conducted.

Embryos were staged as follows: 1-cell stage embryos were collected prior to nuclear envelope breakdown. For 2-cell stage, 1-cell embryos were collected and monitored until they divided. Once the first embryo reached the 2-cell stage, a timer was started, and collection of 2-cell embryos continued for 5 minutes. Similarly, for the 4-cell stage, 2-cell embryos were collected and monitored until they divided. When the first embryo reached the 4-cell stage, a timer was started, and collection of 4-cell embryos continued for 5 minutes. For the 8-cell stage, 1- and 2-cell embryos were collected. When a P1 cell divided, a 3-minute timer was started, 9 and embryos were collected in pools. Each pool was timed for 10 minutes. After washing (taking ∼2-3 mins) we checked that no 4-cell embryos were observed under the microscope prior to freezing. For each stage, this process was repeated until 9 embryos were collected in a single PCR tube.

#### Library Preparation

We used a slightly modified version of the previously described Ribo-ITP protocol ^23^and a detailed protocol can be obtained from (https://ceniklab.github.io/ribo_itp/) which can be referred to for more details. In brief, the PCR tubes with 9 embryos were centrifuged to collect them at the bottom. We then added 2.5µl of 100µg/ml cycloheximide to the PCR tubes and we lysed the embryo by freeze thawing them in liquid nitrogen. We then added 2.5 µl of 2x lysis buffer (40mM Bistris, 100mM NaCl, 2% Triton-X-100, 10mM MgCl_2_, 10 mM CaCl_2_) and mixed well by pipetting. To obtain ribosome protected fragments, we added 1µl of 1:50 diluted MNase and incubated at 37°C for 30 mins. The digestion was then stopped using 1µl of 70mM EGTA and vortexed for at least 30 seconds and stored on ice. The microfluidics chip for ribosome profiling via isotachophoresis was setup and run as provided in ^23^. The ribosome protected fragments were collected in dephosphorylation buffer from the Diagenode D-Plex Small RNA-seq Kit (Diagenode Cat. No C05030001) kit. The replicates for 1-cell (WT), 2-cell (WT), 4-cell (WT), 8-cell (WT), 1-cell (*zu405*), 2-cell (*zu405*) and 4-cell (*zu405*) were two, four, four, five, three, six, and four, respectively. We made a few modifications to the kit’s library preparation protocol, and they were as follows. At the RNA-tailing step we used 0.5µl of Dephosphorylation Reagent along with 0.5µl of T4 Polynucleotide Kinase (New England Biolabs M0201L). Upon adding these two reagents the mixture is incubated for 25 mins at 37°C. Next, during the reverse transcription with template switching, we diluted the TSO by half and used 2µl in the mixture. We used 16 cycles for the final PCR amplification to produce the DNA library. RNA libraries were prepared using the NEBNext® Single Cell/Low Input RNA Library Prep Kit for Illumina® (NEB #E6420S/L), following the manufacturer’s protocol with some modifications. Briefly, embryo samples, each consisting of 9 embryos, were collected in PCR tubes containing 4.5 µl of water. These samples underwent a freeze-thaw cycle using liquid nitrogen to lyse the cells and release the RNA. For library preparation, we employed a two-step PCR amplification process. The initial cDNA enrichment PCR was performed for 14 cycles, followed by a final library amplification PCR for 8 cycles. Three independent biological replicates were prepared for each experimental condition.

#### Fixation and imaging of *C.elegans* embryos

To visualize CAM-1::mNG, LAG-1::mNG and DSH-2::mNG gravid hermaphrodites containing one row of embryos were selected alongside N2 controls. Twenty worms were dissected in 50 µL of 1X egg buffer (5 mM HEPES pH 7.4, 118 mM NaCl, 40 mM KCl, 3.4 mM MgCl_2_, 3.4 mM CaCl_2_) on a watch glass. One-cell embryos were collected within 2 minutes using 2.5 µL of 10% sodium hypochorite solution treatment and cleaned through sequential washes in Shelton’s growth media (SGM; 0.5 mg/mL Inulin,5 mg/mL PVP, 1x BME vitamins, 1x Chemically Defined Lipid Concentrate and 1x Penicillin-Streptomycin added to Schneider’s Drosophila Media) supplemented with 35% fetal bovine serum and then in 1X egg buffer. The following fixation protocol was modified from ^85^. Embryos were fixed in pre-chilled (−20°C) methanol followed by two freeze-thaw cycles in liquid nitrogen. Samples were pelleted at 2000 × g in 30-second intervals with 180° rotation between spins for up to 5 minutes. After methanol removal, samples were incubated in −20°C acetone for 3 minutes, followed by centrifugation for up to 5 minutes. Fixed embryos underwent three 5-minute washes in 1X PBST, followed by incubation in 5% w/v BSA in PBST for 1 hour at 37°C with nutation. Samples were counterstained with DAPI (200 ng/µL) for 10 minutes and washed twice in PBST. Embryos were resuspended in 50 µL PBST and incubated at 4°C for 30 minutes. For imaging, 5 µL of sample was mounted between an L-polylysine-coated coverslip and glass slide, sealed with nail polish, and imaged using a Nikon Ti2 microscope controlled by µManager software and equipped with a 60x, 1.42 NA objective lens; a Visitich iSIM super-resolution confocal scan head; and a Photometrics Kinetix 22 camera. Imaging parameters included 405 nm and 505 nm channels with a 445-505-561-638 dichroic mirror. For live imaging, worms were dissected in egg buffer on L-polylysine-coated coverslips. Samples were sealed onto glass slides using valap and imaged immediately using identical conditions as the fixed samples.

### 13.3. Quantification AND Statistical analysis

#### Read alignment of ribosome profiling and RNA-seq data

Ribosome profiling data were processed using RiboFlow^77^ Input data included gzipped fastq files containing ribosome profiling sequencing data, Bowtie2 index files for the transcriptome reference (appris_celegans_v1) and filter reference (rRNA sequences), a bed file defining CDS, UTR5, and UTR3 regions, and a two-column .tsv file containing transcript lengths. The reference files can be found here^44^ : https://github.com/RiboBase/reference_caenorhabditis-elegans

Unique Molecular Identifiers (UMIs) were extracted from the reads using UMI-tools (Version 1.1.2) with the parameters “-p “^(?P<umi_1>.{12})(?P<discard_1>.{4}).+$” -- extract-method=regex”. Reads were processed using default settings^77^. Alignments with a mapping quality greater than two were retained. Deduplication was performed using UMI-tools^78^. Only footprints between 21 and 40 nucleotides in length were considered for the final analysis.

RiboFlow was also used to process the RNA-seq data. The processing steps were as follows: First, quality control and trimming were conducted using cutadapt (version 4.7), where the first and last 25 nucleotides were trimmed from each read with the parameters ‘-u 25 -u −25 --quality-cutoff=28’. Next, filtered reads were aligned using Bowtie2 in a two-step process: reads were initially aligned to a reference of rRNAs using the parameters ‘-L 15 --no-unal’, and then unaligned reads were aligned to the transcriptome reference using the same parameters. Finally, in the post-alignment processing step, only alignments with a mapping quality greater than 2 were retained for further analysis.

#### Data Filtering and Processing

For each library, we first determined the footprint size range to be used in downstream analyses. Specifically, we selected the ones for which the percentage that mapped to CDS was greater than equal to 92.5 percent. Histone genes were excluded from the analysis due to their lack of poly-A tails, as our RNA-seq was generated using a oligo-dT based reverse transcription strategy (Table S1). A prior study has shown a systematic underestimation of histone transcript levels that prevents reliable calculation of their translational efficiency^86^. Histone genes were identified as the set of genes whose Wormbase Genetic gene nameID begins with “his”. We retained genes with a minimum of 3 counts per million in at least 10 of the 29 collected samples. Batch correction was then performed using ComBat-seq^81^ by providing the day of library prep as the batch variable. These corrected counts were used for all subsequent analyses. We retained genes with a minimum of 10 counts per million in at least 18 of the 21 collected samples. Batch correction was then performed using ComBat-seq^81^ by providing the day of library prep as variable to adjust for potential batch effects. These corrected counts were used for all subsequent analyses. Our final gene set consisted of 4,905 genes that passed both the ribosome profiling and RNA-seq filtering processes.

#### RNA abundance and Ribosome Occupancy correlation analysis

To analyze the relationship between ribosome occupancy and RNA abundance, we performed pairwise correlation analyses using normalized count data. Raw counts were first converted to counts per million (CPM) to account for differences in sequencing depth across samples. For both RNA-seq and Ribo-seq datasets, we calculated the mean CPM values across replicates for each gene. To assess data quality and reproducibility, we computed Spearman correlations between all possible pairs of replicates within each assay type (RNA-seq or Ribo-seq) and took their mean to obtain reliability scores. The observed correlation between mean ribosome occupancy and RNA abundance was then calculated using Spearman’s rank correlation coefficient. To account for measurement noise in both assays, we applied a correction to the observed correlation using the reliability scores, calculated as:

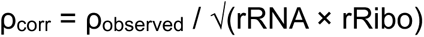

Where rRNA and rRibo represent the reliability scores for RNA-seq and Ribo-seq measurements, respectively. Statistical analyses and visualizations were performed using R.

We analyzed datasets from two independent studies: Lee et al.(2020) and Quarato et al. (2021) to investigate the relationship between mRNA abundance and ribosome occupancy. First, we identified a set of common genes present in our and their datasets. For the Lee et al. (2020) dataset, we extracted average mRNA abundance (measured in Fragments Per Million, FPM) and average ribosome occupancy (also in FPM) for each gene. Similarly, for the Quarato et al. (2021) dataset, we extracted RNA abundance and ribosome occupancy values (measured in Transcripts Per Million, TPM).

#### Pairwise differential expression analysis

Differential expression analysis was performed using the edgeR package (version 4.0.16) in R. Library sizes were normalized using the trimmed mean of M-values method^87^. To identify differentially expressed genes between conditions, quasi-likelihood F-tests were performed using the glmQLFTest function. Contrasts were defined to specify the comparisons of interest, such as differences between consecutive developmental stages or between OMA-1/WT conditions in the case of comparing RNA abundance and ribosome occupancy. Translational efficiency changes were assessed using contrasts that compared the differences between Ribo-seq and RNA-seq data across different conditions or developmental stages

((Condition B Ribo-seq - Condition B RNA-seq) - (Condition A Ribo-seq - Condition A RNA-seq)).

This approach allowed for the identification of genes with significant changes in translational efficiency between the compared conditions, independent of changes in mRNA levels. For each comparison, genes with a false discovery rate (FDR) < 0.2 and |log_2_FC| >1 were considered significantly differentially expressed.

#### Normalization of TE

To get an estimate of TE at a given stage, we calculated the centered log-ratio (CLR) that account for the compositional nature of sequencing data^88^. The CLR of a gene is calculated as the natural logarithm of the ratio between x (defined as the count plus a pseudocount of 1) and the geometric mean of all genes in each replicate and sample.

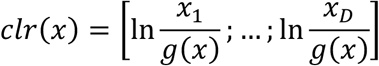

A pseudo count of +1 was added avoid zeros present in the ribosome profiling data. The resulting log-ratios were averaged across replicates. Translational efficiency (TE) was calculated as the difference between the averaged log-ratios of ribosome profiling and total RNA.

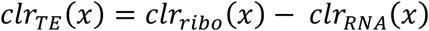

This processed dataset, containing gene names and corresponding TE values, forms the basis for subsequent analyses of translational regulation during early embryonic development in *C. elegans*, both in wild-type and OMA-1 (*zu405*) conditions.

We compared our TE to two prior studies that obtained ribosome profiling data from *C.elegans* embryos^21,22^. For the Quarato et al. dataset^21^, we used their reported centered log-ratio transformed translation efficiency values. For the Lee et al. dataset^22^, we calculated log2 translation efficiency directly from their reported ribosome occupancy and mRNA abundance measurements. To assess the consistency of translation efficiency patterns across studies, we performed a comparative analysis between our stage-specific translation efficiency data and those from both published datasets. We focused on four early developmental stages: 1-cell, 2-cell, 4-cell, and 8-cell embryos. The spearman rank correlation was calculated between these two datasets.

#### Translational efficiency based on localization of transcripts

To categorize genes based on their spatial expression patterns, we utilized differential expression data from Tintori et al.^8^ This dataset compared expression levels in whole embryos to those in germline precursor cells at two-cell, four-cell, and eight-cell stages. Genes were classified into three categories:

1. Somatic genes: log_2_ fold change > 1, adjusted p-value < 0.1, and log_2_ CPM > 4
2. Germline genes: log_2_ fold change < −1, adjusted p-value < 0.1, and log_2_ CPM > 4
3. Evenly expressed genes: |log_2_ fold change| < 1, adjusted p-value > 0.2, and log_2_ CPM > 4

The fold change values represent the expression ratio of (whole embryo)/(germline precursor cell). Thus, positive fold changes indicate higher expression in somatic cells, while negative fold changes suggest enrichment in germline precursors. Statistical differences in translational efficiency between somatic and germline-enriched transcripts were assessed using two-sided Wilcoxon rank-sum tests for each developmental stage. P-values < 0.05 were considered significant and visualized using asterisks (*p < 0.05, **p < 0.01, ***p < 0.001). Summary statistics including median, mean, standard deviation, and quartile values were calculated on R for each group at each developmental stage (see code).

#### TE of P-granule localized transcripts

The localization of transcripts to P-granules was determined using single-molecule fluorescence in situ hybridization data from Scholl et al.^40^. For each transcript, P-granule association was assessed at the 2-cell (P1 cell), 4-cell (P2 cell), and 8-cell (P3 cell) stages. Differential translational efficiency between P-granule localized (“+”) and non-localized (“-”) transcripts was evaluated using two-sided Wilcoxon rank-sum tests at each developmental stage. For transcripts that transitioned into P-granules (from “-” to “+”) between stages, changes in TE were assessed using paired Wilcoxon tests against a null hypothesis of no change. Additionally, differences in TE between Group I transcripts (maintained in P-granules through primordial germ cells) and Group II transcripts (transiently associated) were evaluated using Wilcoxon rank-sum tests at each stage. To assess the enrichment of P-granule transcripts in germline cells, Fisher’s exact tests were performed comparing P-granule-localized transcripts with germline-enriched transcripts at each developmental stage. P-values < 0.05 were considered significant for all statistical tests.

#### Linear regression analysis of RNA-binding proteins

To investigate the relationship between 3’ UTR motifs, RNA-binding protein interactions, and TE, we performed a series of analyses on data from one-cell, two-cell, four-cell, and eight-cell stage embryos. First, we converted the 3’ UTR motif count data for POS-1, MEX-3, and LIN-41 into binary format, where the presence of a motif was denoted as 1 and absence as 0. For OMA-1 and GLD-1, we used binary data from pull-down experiments to indicate protein-RNA interactions. We then merged this binary data with the log-normalized TE data based on gene names. A linear regression model was constructed to assess the influence of various factors on TE. For each developmental stage, we used the following predictors: OMA-1 pull-down status, presence of motifs for POS-1, MEX-3, and LIN-41, and GLD-1 pull-down status. The model for a given stage is represented as:

Translational Efficiency at stage X∼ OMA_pulldown + POS1_motif + MEX3_motif + LIN41_motif + GLD_pulldown

We observed that 815 out of the 1039 transcripts pulled down with OMA-1 in oocytes^34^ were detected robustly in our dataset. For each developmental stage (1-cell, 2-cell, 4-cell, and 8-cell), differences in translational efficiency (TE) between OMA-1-bound (n = 815) and unbound (n = 4090) transcripts were assessed using a two-sided Wilcoxon rank-sum test. The test statistic W is calculated as:

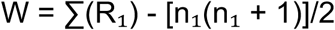

where R₁ is the sum of ranks for OMA-1-bound transcripts, n₁ is the number of bound transcripts, and the second term [n₁(n₁ + 1)]/2 represents the expected sum of ranks under the null hypothesis. The null hypothesis H₀ states that the TE distributions of bound and unbound transcripts are identical, while the alternative hypothesis H₁ states that the TE distributions differ between the two groups. This test was also applied for the violin plots generated for results in Figure S4.

#### Clustering analysis and GO enrichment

For wild-type (WT) data analysis, clustering was performed using k-means algorithm (k=9) on mean-centered data, where the mean expression value of each gene across all stages was subtracted. For visualization, the data was then normalized to the 1-cell stage by subtracting each gene’s 1-cell stage value from all subsequent stages. For the comparative analysis between WT and OMA-1-depleted (OMA) conditions, data was first normalized to the 1-cell stage, then k-means clustering was performed with k=2 clusters for each condition independently on this 1-cell normalized data. The clustering results were combined to create multimodal clusters. Gene ontology (GO) enrichment analysis was conducted using the clusterProfiler package with the org.Ce.eg.db database for *C. elegans*. The background gene set for enrichment analysis was derived from all detected genes in the experiment. GO terms were simplified using the simplify() function with a cutoff of 0.5, and filtered based on enrichment ratio (>2) and adjusted p-value (<0.1). The complete GO enrichment results can be found in the supplement (Table S2 and S5). For the main figure visualization, GO terms were manually curated to remove redundancy and highlight biologically relevant annotations.

