## Supplemental figures and tables for "Landscape and regulation of mRNA translation in the early *C. elegans* embryo"

### Supplementary Figures

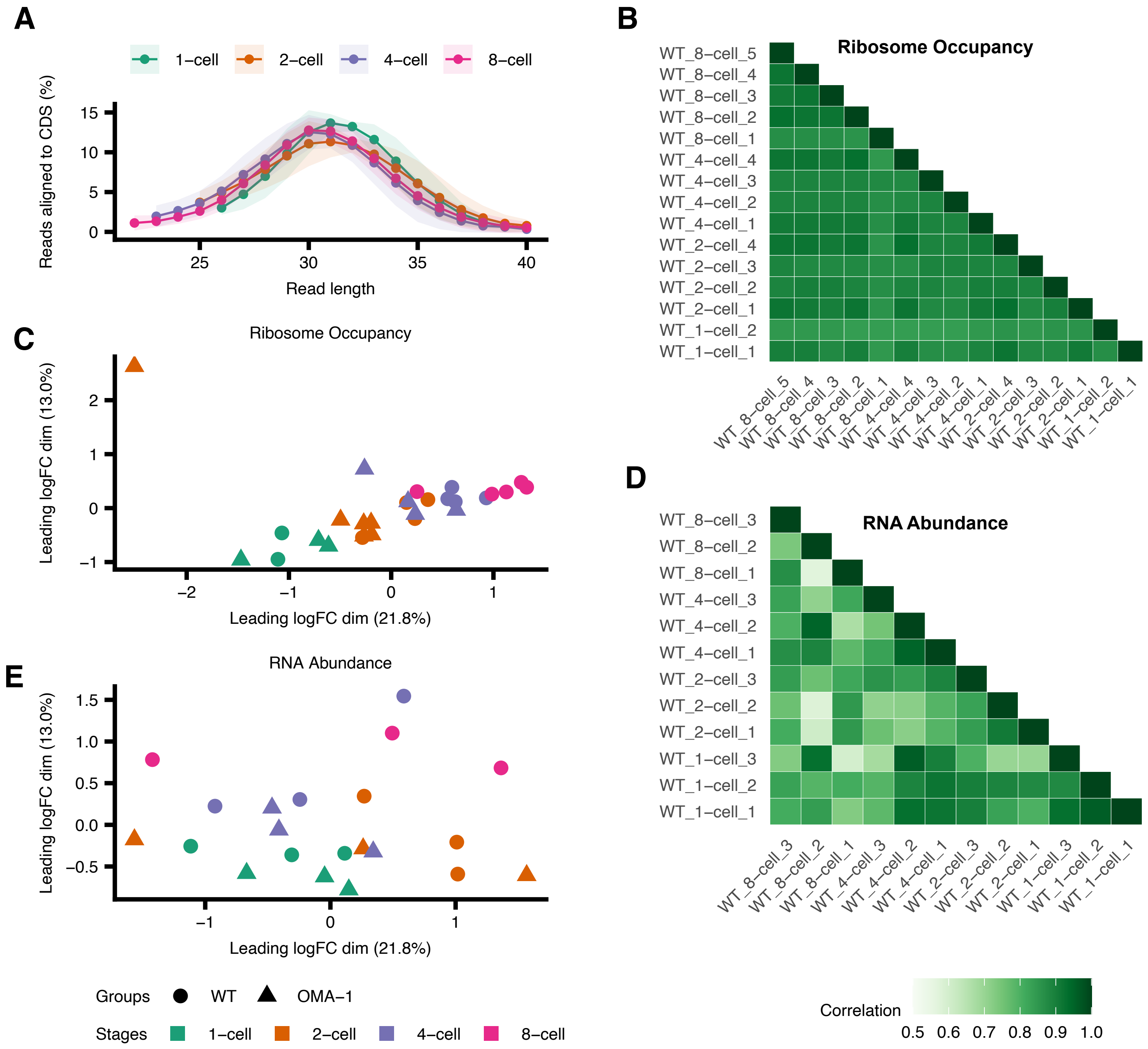
A
1−cell
2−cell
4−cell
8−cell
Reads aligned to CDS (%)
Read length
B
Ribosome Occupancy
WT_8−cell_5
WT_8−cell_4
WT_8−cell_3
WT_8−cell_2
WT_8−cell_1
WT_4−cell_4
WT_4−cell_3
WT_4−cell_2
WT_4−cell_1
WT_2−cell_4
WT_2−cell_3
WT_2−cell_2
WT_2−cell_1
WT_1−cell_2
WT_1−cell_1
C
Ribosome Occupancy
Leading logFC dim (13.0%)
Leading logFC dim (21.8%)
D
RNA Abundance
WT_8−cell_3
WT_8−cell_2
WT_8−cell_1
WT_4−cell_3
WT_4−cell_2
WT_4−cell_1
WT_2−cell_3
WT_2−cell_2
WT_2−cell_1
WT_1−cell_3
WT_1−cell_2
WT_1−cell_1
E
RNA Abundance
Leading logFC dim (13.0%)
Leading logFC dim (21.8%)
Groups WT OMA−1
Stages 1−cell 2−cell 4−cell 8−cell
Correlation
0.5 0.6 0.7 0.8 0.9 1.0

Figure S1:

(A) Distribution of ribosome profiling read lengths across developmental stages, with shaded areas indicating standard deviation between biogical replicates. (B) Correlation matrix of ribosome occupancy, where darker colors indicate stronger correlations. (C) Multidimensional scaling analysis of Ribo-seq. Axes represent arbitrary units. (D) Correlation matrix of RNA abundance between replicates, where darker colors indicate stronger correlations. (E) RNA-seq replicates, with shapes denoting WT versus OMA-1 mutants and colors indicating cell stages. Axes represent arbitrary units.

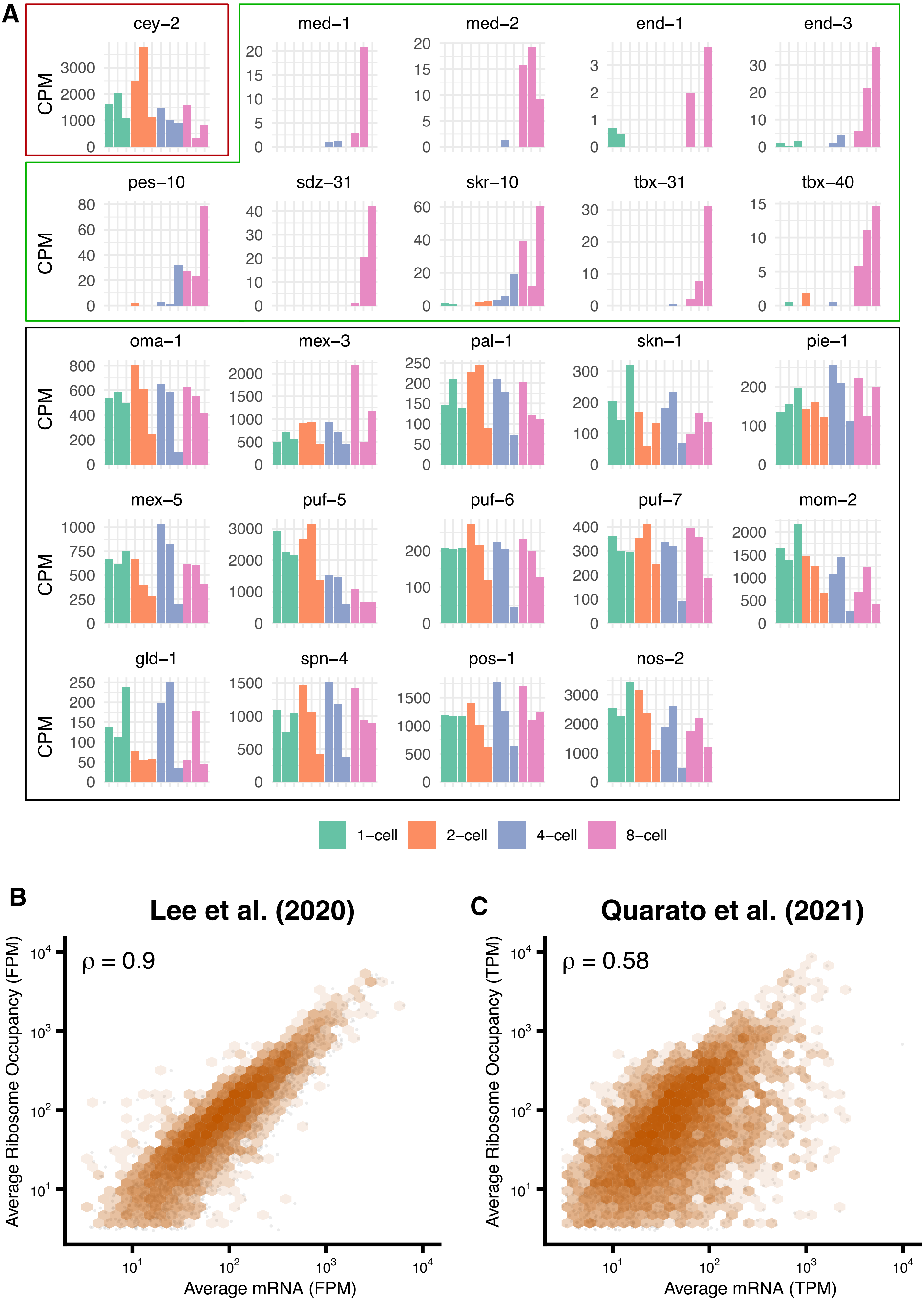

A
cey−2
med−1
med−2
end−1
end−3
pes−10
sdz−31
skr−10
tbx−31
tbx−40
oma−1
mex−3
pal−1
skn−1
pie−1
mex−5
puf−5
puf−6
puf−7
mom−2
gld−1
spn−4
pos−1
nos−2
CPM
1−cell
2−cell
4−cell
8−cell
B
Lee et al. (2020)
ρ = 0.9
Average Ribosome Occupancy (FPM)
Average mRNA (FPM)
C
Quarato et al. (2021)
ρ = 0.58
Average Ribosome Occupancy (TPM)
Average mRNA (TPM)

Figure S2:

(A) Expression patterns from RNA-seq of marker genes, highlighting cey-2 in red box showing progressive degradation throughout embryogenesis, while genes predominantly expressed at the 8-cell stage are in green box. Other previously identified maternally deposited mRNA are shown in the black box. Each bar in the plot of the same color represents a replicate. Relationship between mRNA abundance and ribosome occupancy from (B) Lee et al. (2022) (C) Quarato et al. (2021). The plot shows average mRNA levels versus average ribosome occupancy on log10 scales. Gray dots represent individual genes with hexagonal binning (orange) indicating data density. The Spearman correlation coefficient (rho) quantifies the relationship strength

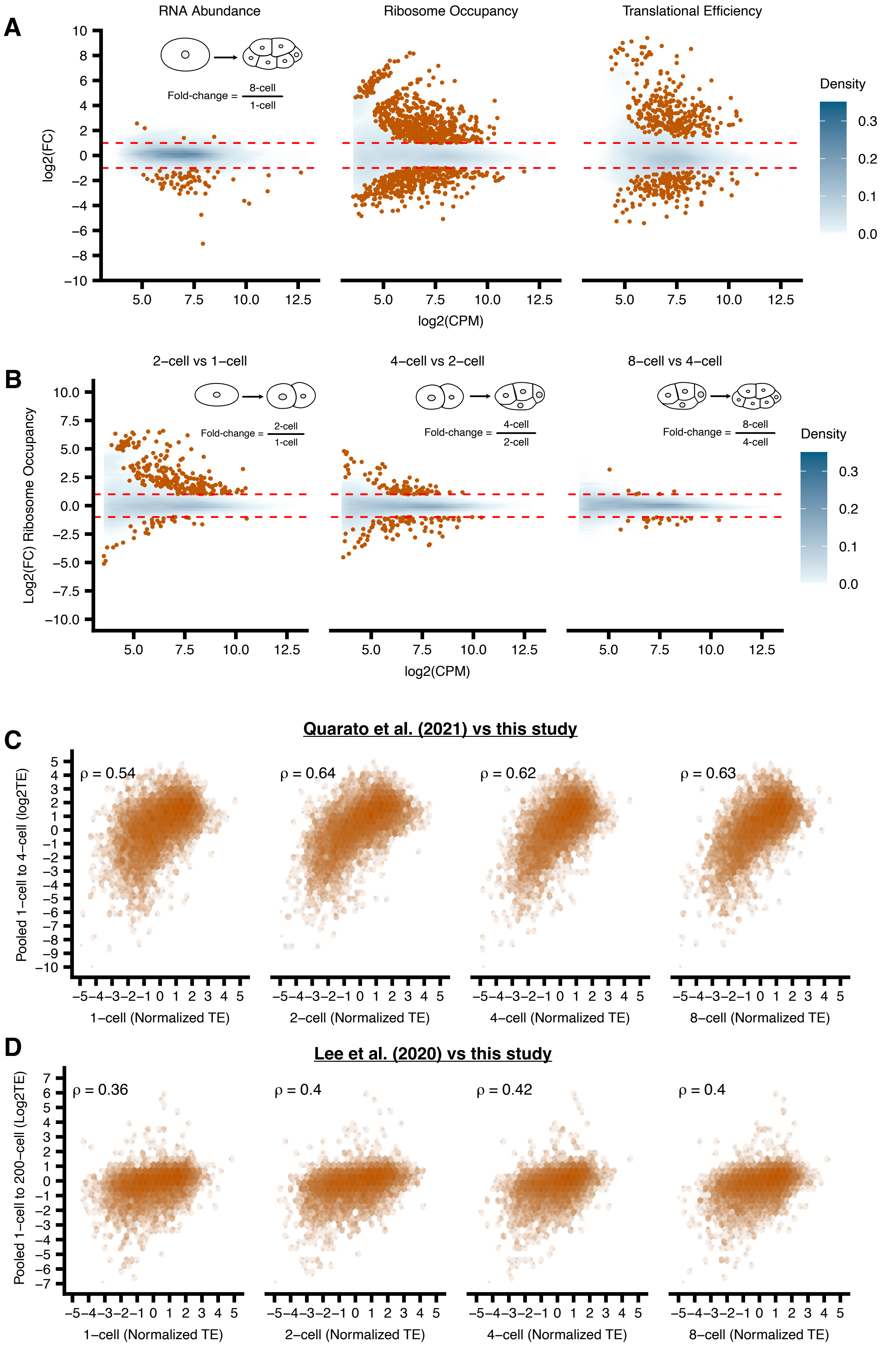

A
RNA Abundance
Ribosome Occupancy
Translational Efficiency
Fold-change = 8-cell / 1-cell
log2(FC)
log2(CPM)
Density
B
2–cell vs 1–cell
4–cell vs 2–cell
8–cell vs 4–cell
Fold-change = 2-cell / 1-cell
Fold-change = 4-cell / 2-cell
Fold-change = 8-cell / 4-cell
Log2(FC) Ribosome Occupancy
log2(CPM)
Density
C
Quarato et al. (2021) vs this study
Pooled 1–cell to 4–cell (log2TE)
ρ = 0.54
ρ = 0.64
ρ = 0.62
ρ = 0.63
1–cell (Normalized TE)
2–cell (Normalized TE)
4–cell (Normalized TE)
8–cell (Normalized TE)
D
Lee et al. (2020) vs this study
Pooled 1–cell to 200–cell (Log2TE)
ρ = 0.36
ρ = 0.4
ρ = 0.42
ρ = 0.4
1–cell (Normalized TE)
2–cell (Normalized TE)
4–cell (Normalized TE)
8–cell (Normalized TE)

Figure S3:

(A) Mean difference plots comparing RNA abundance, ribosome occupancy, and translational efficiency between 8-cell and 1-cell stage wild-type embryos. X-axis shows expression level (log2CPM), Y-axis shows $\log_2$ fold change. Mean difference plots comparing gene expression between sequential *C. elegans* embryonic stages (B)1-cell to 2-cell, 2-cell to 4-cell, and 4-cell to 8-cell. The log2 fold change in ribosome occupancy (y-axis) is plotted against the mean of normalized counts (x-axis). Blue density plot represents the overall distribution of genes with the intensity corresponds to the density of points. The orange points indicate transcripts with significant expression changes (FDR < 0.2 and $\log_2$FC >1 or < -1). Blue density indicates concentration of data points. Orange points highlight significant changes ($|\log_2 FC| > 1$, FDR < 0.05). Red dashed lines indicate ±1 $\log_2$FC thresholds. Correlations between stage-specific and pooled embryonic translational efficiencies. (C) Comparison of normalized TE at the same developmental stages with pooled 1-cell to 4-cell TE data from Quarato et al. (2021) (y-axes). (D) Comparison of normalized translational efficiency (TE) at 1-cell, 2-cell, 4-cell, and 8-cell stages (x-axes) with pooled 1-cell to 200-cell TE data from Lee et al. (2020) (y-axes). Orange hexagonal binning shows data density with Spearman correlation coefficients (rho) indicating relationship strength.

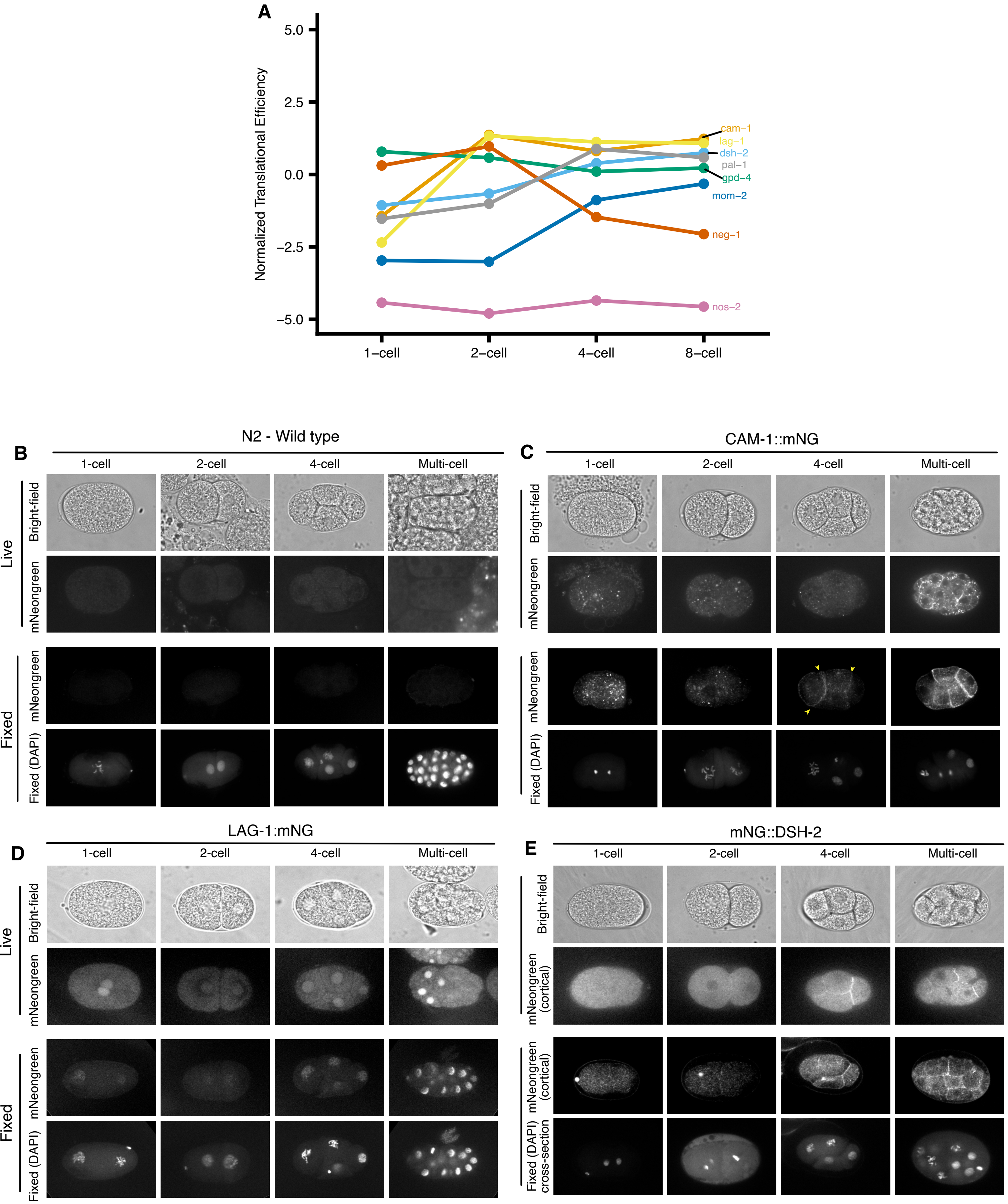
A
Normalized Translational Efficiency
5.0
2.5
0.0
−2.5
−5.0
1−cell
2−cell
4−cell
8−cell
cam−1
lag−1
dsh−2
pal−1
gpd−4
mom−2
neg−1
nos−2
B
N2 - Wild type
C
CAM-1::mNG
D
LAG-1:mNG
E
mNG::DSH-2
1-cell
2-cell
4-cell
Multi-cell
Live
Fixed
Bright-field
mNeongreen
mNeongreen (cortical)
Fixed (DAPI)
Fixed (DAPI) cross-section

Figure S4:

(A) Normalized TE profiles of previously identified (PAL-1, MOM-2, NEG-1, NOS-2) and newly identified (CAM-1, LAG-1 and DSH-1) translationally regulated transcripts compared to the housekeeping gene GPD-4. (B-E) Localization of newly identified proteins during early embryonic development. (B) N2 control, (C) CAM-1::mNG, (D) LAG-1::mNG, and (E) DSH-1::mNG embryos imaged at 1-cell, 2-cell, 4-cell, and multicellular stages. For each genotype: brightfield (top row), live mNG fluorescence (middle row), and fixed samples showing mNG signal with DAPI counterstain (bottom row). Arrowheads indicate cell-cell contact localization of fluorescently tagged proteins in CAM-1::mNG.

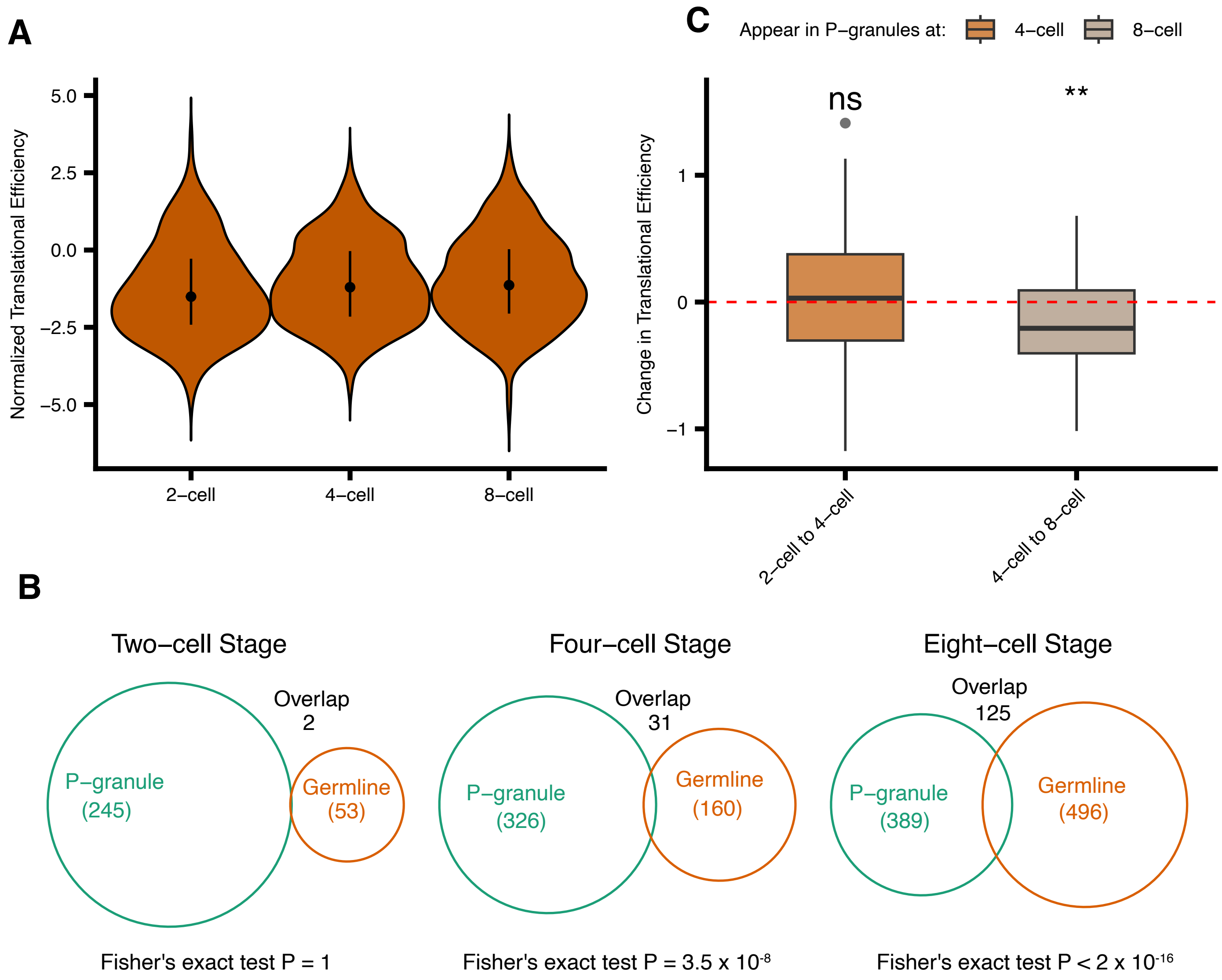
A
Normalized Translational Efficiency
5.0
2.5
0.0
−2.5
−5.0
2−cell
4−cell
8−cell
C
Appear in P−granules at:
4−cell
8−cell
ns
**
Change in Translational Efficiency
1
0
−1
2−cell to 4−cell
4−cell to 8−cell
B
Two−cell Stage
Overlap
2
P−granule
(245)
Germline
(53)
Fisher's exact test P = 1
Four−cell Stage
Overlap
31
P−granule
(326)
Germline
(160)
Fisher's exact test P = 3.5 x 10-8
Eight−cell Stage
Overlap
125
P−granule
(389)
Germline
(496)
Fisher's exact test P < 2 x 10-16

Figure S5:

(A) Translational efficiency of P-granule localized transcripts across developmental stages. Violin plots show distribution of TE values for P-granule transcripts at 2-cell (n=245), 4-cell (n=326), and 8-cell (n=389) stages, with median and quartiles indicated. (B) The overlap between P-granule-associated genes and genes expressed at specific developmental stages in *C. elegans* early embryogenesis. The Venn diagrams illustrate three developmental timepoints: two-cell, four-cell, and eight-cell stage. (C) Changes in translational efficiency of transcripts as they transition into P-granules. Box plots show TE changes for transcripts first appearing in P-granules at 4-cell (orange, n=81) and 8-cell stages (brown, n=43). Red dashed line indicates no change in TE. Asterisks denote significant differences from zero (***p < 0.001, Wilcoxon test).

**A**

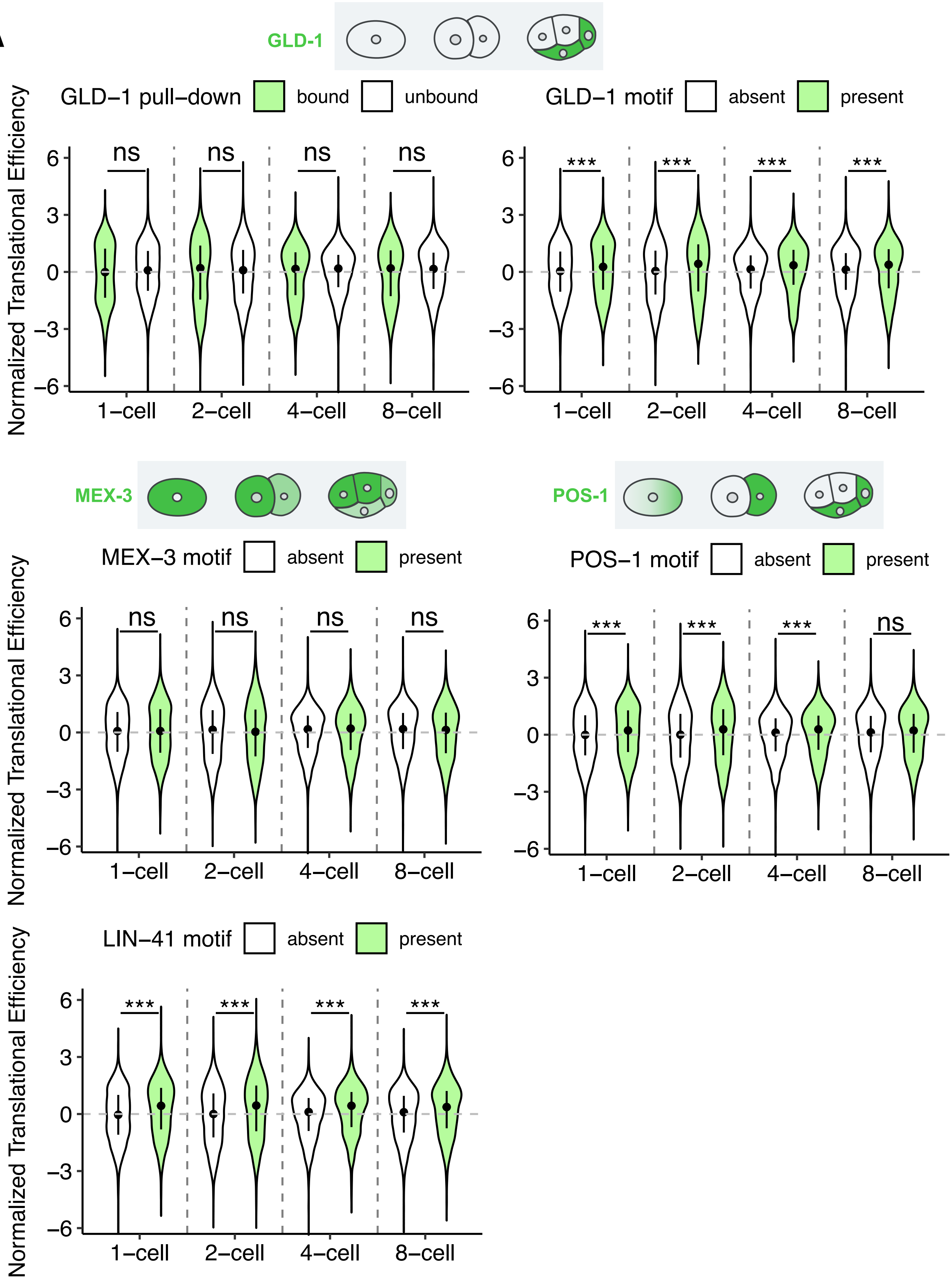
GLD-1
GLD−1 pull−down
bound
unbound
GLD−1 motif
absent
present
Normalized Translational Efficiency
ns
***
1−cell
2−cell
4−cell
8−cell
MEX-3
POS-1
MEX−3 motif
POS−1 motif
LIN−41 motif

Figure S6:

Violin plots showing the distribution of normalized translational efficiency for transcripts across early *C. elegans* embryonic stages (1-cell, 2-cell, 4-cell, and 8-cell). Comparisons are shown between: (A) GLD-1-bound versus unbound transcripts from RNA-immunoprecipitation data, and transcripts with present versus absent motifs for (B) POS-1, (C) MEX-3, (D) GLD-1, and (E) LIN-41. Green violins represent bound/motif-present transcripts, while white violins indicate unbound/motif-absent transcripts. Plot width indicates probability density of translational efficiency values(***$p < 0.001$, Wilcoxon test). Illustrations indicate known localization of proteins during early embryogenesis

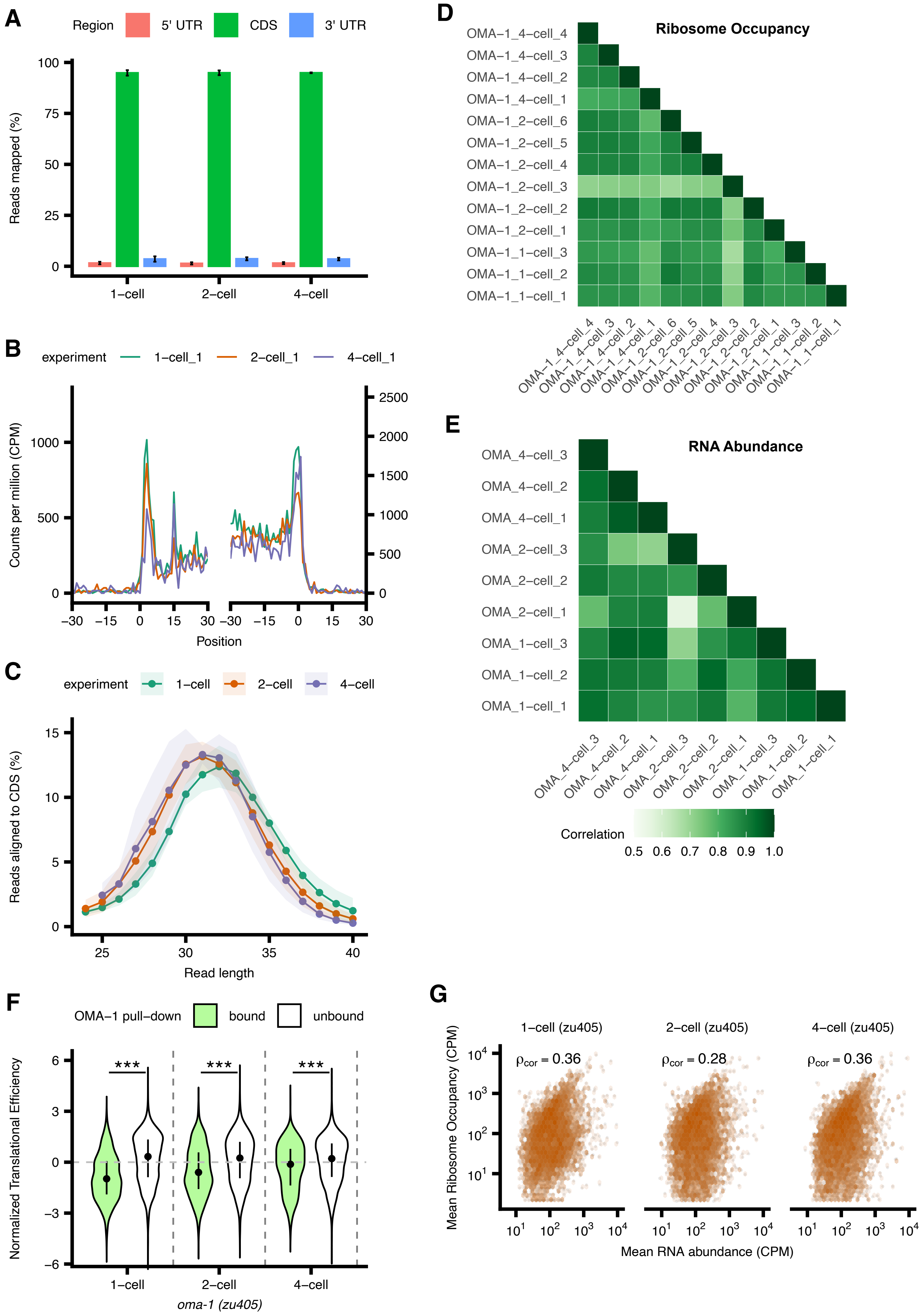

A
Region
5' UTR
CDS
3' UTR
Reads mapped (%)
1−cell
2−cell
4−cell
B
experiment
1−cell_1
2−cell_1
4−cell_1
Counts per million (CPM)
Position
C
experiment
1−cell
2−cell
4−cell
Reads aligned to CDS (%)
Read length
D
Ribosome Occupancy
OMA−1_4−cell_4
OMA−1_4−cell_3
OMA−1_4−cell_2
OMA−1_4−cell_1
OMA−1_2−cell_6
OMA−1_2−cell_5
OMA−1_2−cell_4
OMA−1_2−cell_3
OMA−1_2−cell_2
OMA−1_2−cell_1
OMA−1_1−cell_3
OMA−1_1−cell_2
OMA−1_1−cell_1
E
RNA Abundance
OMA_4−cell_3
OMA_4−cell_2
OMA_4−cell_1
OMA_2−cell_3
OMA_2−cell_2
OMA_2−cell_1
OMA_1−cell_3
OMA_1−cell_2
OMA_1−cell_1
Correlation
0.5 0.6 0.7 0.8 0.9 1.0
F
OMA−1 pull−down
bound
unbound
Normalized Translational Efficiency
***
1−cell
2−cell
4−cell
oma-1 (zu405)
G
1−cell (zu405)
2−cell (zu405)
4−cell (zu405)
ρcor = 0.36
ρcor = 0.28
ρcor = 0.36
Mean Ribosome Occupancy (CPM)
Mean RNA abundance (CPM)

Figure S7:

(A)The allocation of ribosome profiling reads across genomic features (CDS, 5' UTR, 3' UTR) is presented for each stage in *oma-1(zu405)* mutants. (B) Ribosome occupancy around the translation start and stop sites in a representative 1-cell, 2-cell and 4-cell *zu405* staged embryo. Translation start (or stop) sites are denoted by the position 0. Aggregated read counts (*y* axis) relative to the start (or stop) sites are plotted after A-site correction (C) Distribution of ribosome profiling read lengths across developmental stages, with shaded areas indicating standard deviation between biological replicates. (D) Correlation matrix of ribosome occupancy and (E) RNA abundance between replicates, where darker colors indicate stronger correlations. (F) Violin plots in illustrate the distribution of normalized translational efficiency for OMA-1-bound (green) and unbound (white) transcripts across three early *oma-1(zu405)* embryonic stages (1-cell, 2-cell and 4-cell) identified from previous RNA-immunoprecipitation data. The width of each violin represents the probability density of efficiency values(***p < 0.001, Wilcoxon test). (G) Pairwise correlation between ribosome occupancy and RNA abundance at the three stages of early *zu405* embryo development is presented. The mean cpm of ribosome occupancy is plot against the mean cpm of RNA abundance at each stage. Rho (cor) is the corrected spearman correlation based on the reliability r (RNA) and r (Ribo) which are the replicate-to-replicate correlation (methods).

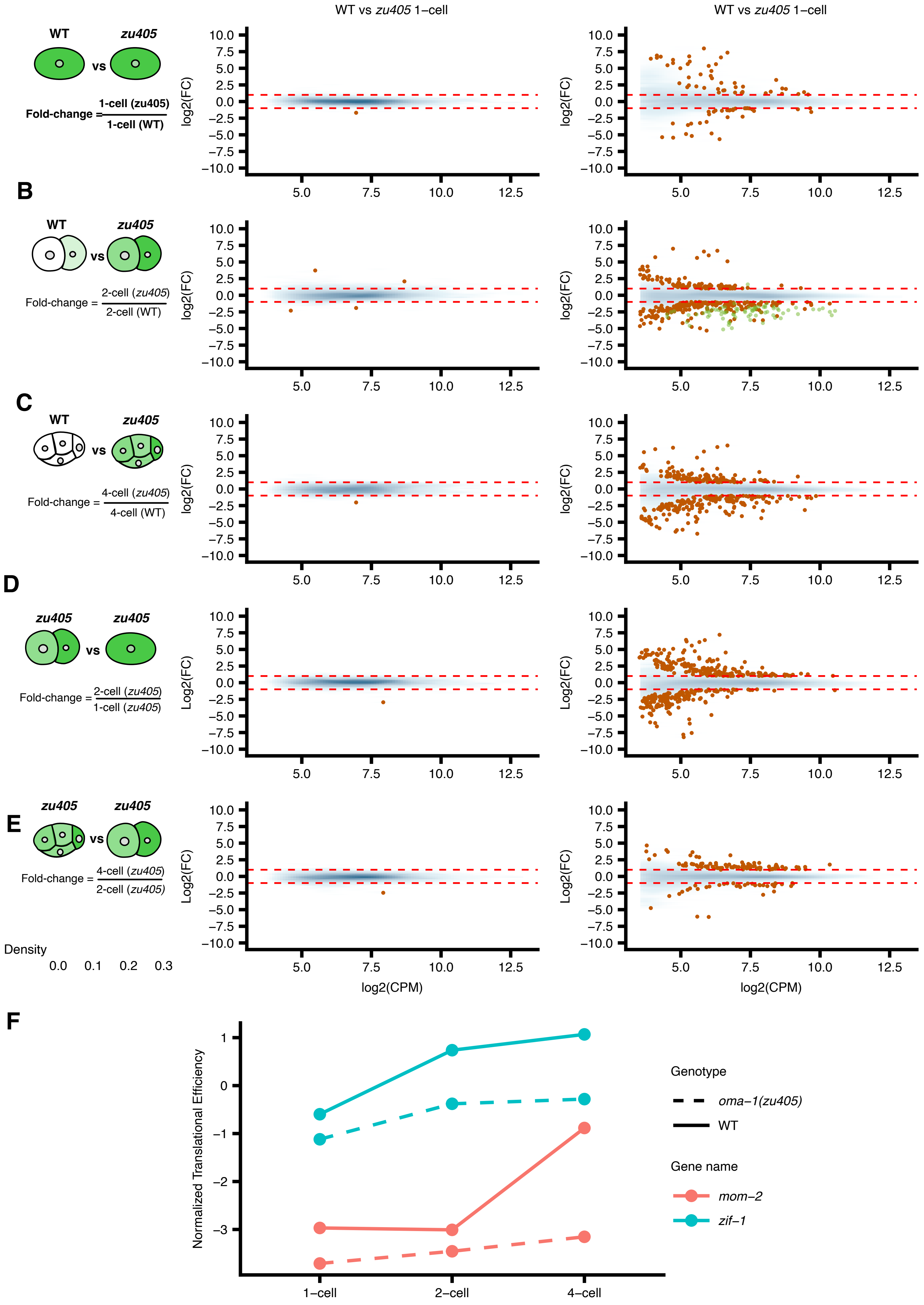

WT vs zu405 1−cell
WT vs zu405 1−cell
WT
zu405
vs
Fold-change = 1-cell (zu405) / 1-cell (WT)
B
WT
zu405
vs
Fold-change = 2-cell (zu405) / 2-cell (WT)
C
WT
zu405
vs
Fold-change = 4-cell (zu405) / 4-cell (WT)
D
zu405
zu405
vs
Fold-change = 2-cell (zu405) / 1-cell (zu405)
E
zu405
zu405
vs
Fold-change = 4-cell (zu405) / 2-cell (zu405)
Density
0.0 0.1 0.2 0.3
log2(FC)
Log2(FC)
log2(CPM)
F
Normalized Translational Efficiency
1−cell
2−cell
4−cell
Genotype
oma−1(zu405)
WT
Gene name
mom−2
zif−1

Figure S8:

Mean difference plots comparing gene expression between *oma-1(zu405)* mutant and wild-type embryos. (A-C) Comparisons between *oma-1(zu405)* and wild-type at 1-cell, 2-cell, and 4-cell stages, respectively. (D-E) Stage-to-stage comparisons within zu405 embryos from 1-cell to 2-cell and 2-cell to 4-cell. For all plots, $\log_2$ fold changes in RNA abundance and translational efficiency (y-axis) are plotted against mean normalized counts (x-axis). Blue density shading indicates the distribution of all genes, with darker shading representing higher point density. Orange points highlight significantly differentially expressed genes (FDR $< 0.2$ and $|\log_2 FC| > 1$). (F) Developmental dynamics of translational efficiency for mom-2 and zif-1 in wild-type and oma-1 mutant embryos. The plot compares normalized translational efficiency (TE) of mom-2 and zif-1 genes across early embryonic stages (1-cell, 2-cell, and 4-cell) in both wild-type (WT, solid lines) and *oma-1(zu405)* mutant (dashed lines) backgrounds.

**Table S1 Count Data for RNA Sequencing and Ribosome Profiling**

Count data for 20,189 transcripts measured by RNA sequencing and ribosome profiling across early embryonic development stages (1-cell, 2-cell, 4-cell, and 8-cell) in both wild-type (WT) and *oma-1 (zu405)* genotypes. Sample names indicate genotype, developmental stage, batch (B#), replicate number, and data type (RNA or RIBO).

**Table S2: Gene Ontology Enrichment Analysis Results for WT**

Results from gene ontology enrichment analysis showing clusters of genes with shared biological functions. The table includes ontology categories, term IDs, descriptions, gene ratios, background ratios, p-values, adjusted p-values, q-values, gene IDs, and count information for 1,726 enriched terms.

**Table S3: Fisher's Exact Test Analysis of P-Granule and Germline-Enriched Gene Overlap**

Statistical analysis of the overlap between P-granule transcripts and germline-enriched genes at three early embryonic stages (two-cell, four-cell, and eight-cell) in C. elegans. The table includes odds ratios, p-values, confidence intervals, and gene counts to quantify the association between P-granule localization and germline enrichment.

**Table S4: Linear Regression Analysis of RNA-Binding Protein Effects on Translation Efficiency**

Results of linear regression analysis showing how translation efficiency correlates with RNA-binding protein interactions (OMA-1, POS-1, MEX-3, GLD-1, and LIN-41) across one-cell, two-cell, and four-cell embryonic stages in C. elegans. The table includes regression coefficients and p-values to quantify how each protein interaction affects translational regulation during early development.

**Table S5: Gene Ontology Enrichment Analysis of OMA-1-Regulated Transcript Clusters**

Gene ontology enrichment analysis results for 127 biological terms associated with the four temporal clusters of OMA-1-bound transcripts shown in Figure 7B. The table provides the ontology categories, term descriptions, gene ratios, enrichment ratios, and statistical significance values (p-values, adjusted p-values, q-values) for functional annotations enriched in each cluster.
